# Hybrid sequencing reveals the genome of a *Chrysochromulina parva* virus and highlight its distinct replication strategy

**DOI:** 10.1101/2025.01.22.633263

**Authors:** Delaney Nash, Christine N. Palermo, Ichiro Inamoto, Trevor C. Charles, Jozef I. Nissimov, Steven M. Short

## Abstract

*Chrysochromulina parva* (*C. parva*) is a eukaryotic freshwater haptophyte algae found in lakes and rivers worldwide. It is known to be infected by viruses, yet knowledge of the diversity and activity of these viruses is still very limited. Based on sequences of PCR-amplified *polB* gene fragments, *Chrysochromulina parva* virus BQ1 (CpV-BQ1) was the first known lytic agent of *C. parva*, and was considered a member of the virus family *Phycodnaviridae,* order *Algavirales*. However, the genome of a different *C. parva*-infecting virus (CpV-BQ2, or *Tethysvirus ontarioense*) from another virus family, the *Mesomimiviridae*, order *Imitervirales,* was the first sequenced. Here, we report the complete genome sequence of the putative phycodnavirus CpV-BQ1, accession PQ783904. The complete CpV-BQ1 genome sequence is 165,454 bp with a GC content of 32.32% and it encodes 193 open reading frames. Phylogenetic analyses of several virus hallmark genes including the DNA polymerase (*polB*), the late gene transcription factor (VLTF-3), and the putative A32-like virion packaging ATPase (Viral A32) all demonstrate that CpV-BQ1 is most closely related to other viruses in the phylum *Megaviricetes* within the order *Algavirales*, family *Phycodnaviridae*.

## Introduction

Research into viruses infecting aquatic microbes has seen a sharp increase in recent years. Many aquatic viruses are Nucleocytoplasmic large DNA viruses (NCLDVs), a diverse group of eukaryotic viruses (1) that were recently classified under the viral phylum *Nucleocytoviricota.* The *Nucleocytoviricota* includes the class *Megaviricetes* with three orders and numerous families including the *Mesomimiviridae* and the *Phycodnaviridae* (2). Viruses belonging to the class *Megaviricetes* are characterized by large double-stranded DNA genomes, typically over 100 kb in size, an ability to replicate in both the host’s nucleus and cytoplasm, and the presence of a core set of shared genes, including those encoding the major capsid protein, DNA polymerase B, and others (3). Viruses in the *Megaviricetes* are also notable for their complexity and size, with some members known as “giant viruses” having genomes and virions that rival those of small cellular organisms (4). They infect a wide range of eukaryotic hosts, from algae to animals, and are found in various environments, particularly aquatic ecosystems (4). They encode genes for DNA replication, transcription, and repair (3), have an extensive genomic plasticity and diversity (3), have acquired genes through lateral gene transfer from various sources (4), and have the potential to significantly alter host metabolism during infection (4).

In addition to their unique genomic characteristics, the *Megaviricetes* play significant ecological roles in various ecosystems, particularly in aquatic environments. They can affect the population dynamics of their unicellular hosts, including controlling algal blooms. For example, *Heterosigma akashiwo* virus (HaV) influences seasonal harmful algal blooms in coastal areas (5). Moreover, viruses in the *Megaviricetes* contribute to biological carbon export from marine surfaces to deep layers through host-cell death (6); can impact nitrogen metabolism and fermentation processes in their hosts (7); and can alter eukaryotic community structures, particularly in marine environments, by infecting various eukaryotic lineages (6,7). Many of these viruses are involved in metabolic reprogramming by encoding genes involved in nutrient uptake, light harvesting, and central carbon metabolism, allowing them to reprogram host metabolism during infection (4). The latter includes genes for photosynthesis, diverse substrate transport, and light-driven proton pumps (1). Indeed, by infecting and lysing their hosts, many *Megaviricetes* drive the release of organic matter into the environment, contributing to nutrient cycling in ecosystems (6,7). Collectively, these ecological roles highlight the importance of *Megaviricetes* in shaping ecosystem dynamics, particularly in aquatic environments, and their potential impact on global biogeochemical cycles.

One of the most understudied aquatic microbial eukaryotes that is infected by viruses that are likely to be classified within the *Megaviricetes* is *Chrysochromulina parva* (*C. parva*), a freshwater haptophyte alga (8,9). It is a small unicellular organism, typically 4-6 μm in size (10) with two flagella (∼8 μm long) and a long haptonema (up to 10 times the body length) (10). *C. parva* has a deep groove running the length of the cell, from which the flagella and haptonema emerge, contains two chloroplasts with internal pyrenoids, each associated with a large lipid body, and has a simple cellular morphology with a eukaryotic nucleus, mitochondria with tubular cristae, and a Golgi apparatus. Unlike most *Chrysochromulina* species, *C. parva* lacks visible scales on its cell surface or within the Golgi apparatus (10).

Viruses capable of infecting and lysing *C. parva* were first isolated in 2015 from a Lake Ontario water sample. This sample, collected from the Bay of Quinte, contained a filterable, heat labile, chloroform sensitive lytic agent that was capable of completely lysing cultures of *C. parva* (11). PCR primers designed to amplify gene fragments from a range of viruses within the *Phycodnaviridae* were used to amplify DNA polymerase B (polB) and major capsid protein (MCP) gene fragments (12). Sanger sequencing of the resulting amplicons led to the identification of a single DNA polB gene fragment and 9 unique MCP fragments. Based on the *polB* phylogeny, the lytic agent was named *Chrysochromulina parva* virus BQ1 (CpV-BQ1) and was putatively classified as a Phycodnavirus; the presence of multiple MCP fragments allowed the researchers to speculate that more than one type of virus might have been isolated from this water sample (11). Further culturing, isolation, and high-throughput sequencing experiments led to the assembly of a giant, 437 kb, virus genome encoding 503 ORFs, as well the ∼23 kb genomes of three polinton-like viruses (PLVs) which are viruses similar to virophages and which need a helper virus for their replication. However, the CpV-BQ1 *polB* gene fragment initially recovered by PCR was not found within this giant virus genome or the PLV genomes, indicating that an additional virus of *C. parva*, named CpV-BQ2, was present in the Bay of Quinte water sample and had been co-cultured along with CpV-BQ1 (13). Here we report the second genome sequence of a *C. parva* virus noting that this genome assembly was derived from CpV-BQ1 because it encodes an identical *polB* gene as that which originally led to the classification of CpV-BQ1 as a member of the *Phycodnaviridae.* In order to study this virus’s genome characteristics, we implemented a two step approach for its sequencing, which combined short read Illumina and long read Nanopore sequencing, coupled with PCR-enabled molecular analysis.

## Materials and Methods

### Host and Virus Cultivation

Viruses infecting *Chrysochromulina parva* strain CCMP 291 (National Center for Marine Algae and Microbiota, East Boothbay, Maine, USA) were originally isolated from an embayment of Lake Ontario, Canada, in 2011 (11) and have been regularly propagated in the laboratory since then. The CpV-BQ1 virus was purified from cultures of mixed *C. parva* viruses, as described in Stough et al. (2019), via an end-point dilution approach. Briefly, serial 10-fold dilutions of *C. parva* lysates were inoculated into 96-well microtiter plates with mid-log phase *C. parva* cells at a concentration of approximately 6.0 x 10^5^ cells/mL as determined using a hemocytometer and light microscope. Medium from individual wells which lysed at the highest dilution level (i.e., lowest concentration) of viruses were transferred in into 50 mL cultures of mid-log *C. parva* and the resulting lysates were filtered through sterile 0.22 µm pore-size PVDF Durapore® membrane filters (EMD Millipore, GVWP00010). This process was repeated 3 times to ensure that only a single type of CpV was propagated, and the presence of CpV-BQ1 was confirmed throughout this purification process using the qPCR method described in Mirza et al. (2015).

### Nucleic Acid Extraction

Following purification of CpV-BQ1, genomic material was prepared for sequencing by extracting nucleic acids from a 600 mL lysate of a *C. parva* culture infected with CpV-BQ1. This lysate was filtered through a 0.22 µm pore-size Steritop® disposable bottle top filters (MilliporeSigma, S2GPT10RE) and was concentrated via ultracentrifugation at 31,000 rpm for 1 hour at 20°C in a SW32Ti rotor (Beckman Coulter Life Sciences). The supernatant was decanted and pelleted material was resuspended in 10 mM Tris-Cl, pH 8.5. Nucleic acids were extracted from the concentrated viral lysate using a Maxwell® RSC Viral Total Nucleic Acid extraction kit (Promega, AS1330) following the manufacturer’s protocol for 300 µL of sample input. The DNA concentration measured using a Qubit fluorometer with a dsDNA HS kit (Thermo Fisher Scientific, Q32851) was 26.2 ng/µL.

### Illumina and Nanopore library preparation and sequencing

The DNA was sent to SeqCenter (Pittsburgh, USA) for paired-end whole genome shotgun sequencing. Sample libraries were prepared using the Illumina DNA Prep kit and IDT 10 base pair UDI indices, and were sequenced on an Illumina NovaSeq 6000, producing 2 x 151 base pair reads. Demultiplexing, quality control, and adapter trimming was performed with BCL-Convert v4.0.3, generating 14,757,009 read pairs.

The DNA was also prepared for Nanopore sequencing at the University of Waterloo. Extracted DNA was diluted to a concentration of 5 ng/µL and prepared with the Nanopore Rapid PCR barcoding 24 V14 kit (Nanopore, SQK-RPB114.24). The prepared DNA library was sequenced on a MinION Mk1B device using an R10.4.1 flow cell (Nanopore, FLO-MIN114).

### Read processing and Genome Assembly

Nanopore sequencing produced a total of 2,325,374 raw reads. First, Filtlong was used to remove reads less than 1000 bp long and 10% of the worst quality reads, resulting in 1,879,244 reads (14). Kraken2 v2.0.7-beta was then used to filter out classified reads, removing contaminating host, bacterial, and human reads. The standard Kraken2 database v9/26/2022 was used with a 0.001 confidence filter and the unclassified-out flag to generate a separate file of unclassified reads, from which 1,048,312 reads were obtained (15).

Filtered long reads and Illumina reads were then used for assembly with the TryCycler pipeline v0.5.4 (16). Briefly, Filtlong was used to keep the best 80% of filtered nanopore reads. This final read filtering step yielded 796,152 reads with an N50 of 6,479bp and a mean length of 6,352 bp (14). Long reads were then subsampled to create 24 read subsets with approximately 8,645 reads per subset. An estimated genome size of 400,000 bp was used to subset reads, which was based on the size of the CpV-BQ2 virus genome (Accession MH918795). Each read subset was assembled with either Flye v2.9.2-b1786, Miniasm & Minipolish v0.1.2, Raven v1.8.1, or Canu v2.2 (17–20). Of the 24 read subsets, 6 were assembled into contigs by each respective tool (17–20). From these 24 genome assemblies, 26 contigs were produced.

Assembled contigs were clustered to determine the MASH distance between assemblies. The MASH distance is based on the Jaccard index and provides a measure of similarity and diversity between each sample. Using MASH distances, a phylogenetic tree was generated during the TryCycler clustering step (Figure 1). Of the 26 assembled contigs, 23 clustered closely together. These assembled contigs were reconciled which ensures contigs are on the same strand and sufficiently similar for downstream assembly. To ensure contigs were sufficiently similar, the TryCycler default thresholds were used, including a 98% sequence identity score and minimum 1-kbp identity score of 25%. Based on these thresholds, 7 of the 23 contigs were discarded. A dotplot was generated to visualize the topology of the remaining 16 contigs (Figure 2).

**Figure 1.**
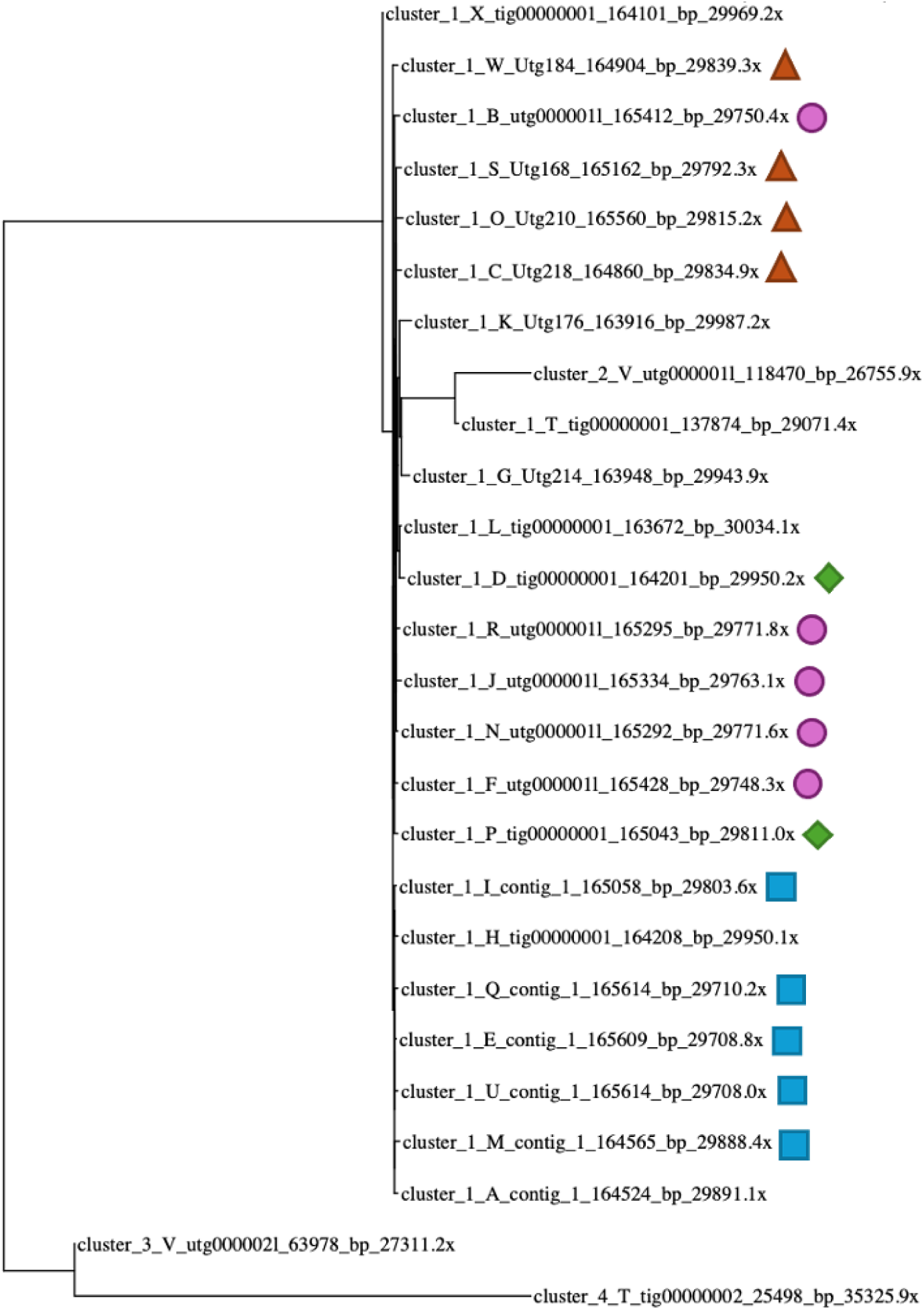
Phylogenetic tree of Clustered Assembled Contigs. The TryCycler clustered assemblies based on the MASH distances between contigs displayed in a phylogenetic tree. Almost all assemblies fall within the cluster 1 branching structure. The 16 assemblies used for reconciling and MSA steps are labeled (e.g., blue squares are Flye assemblies, pink circles are Minipolish assemblies, orange triangles are Raven assemblies, and green diamonds are Canu assemblies).

**Figure 2.**
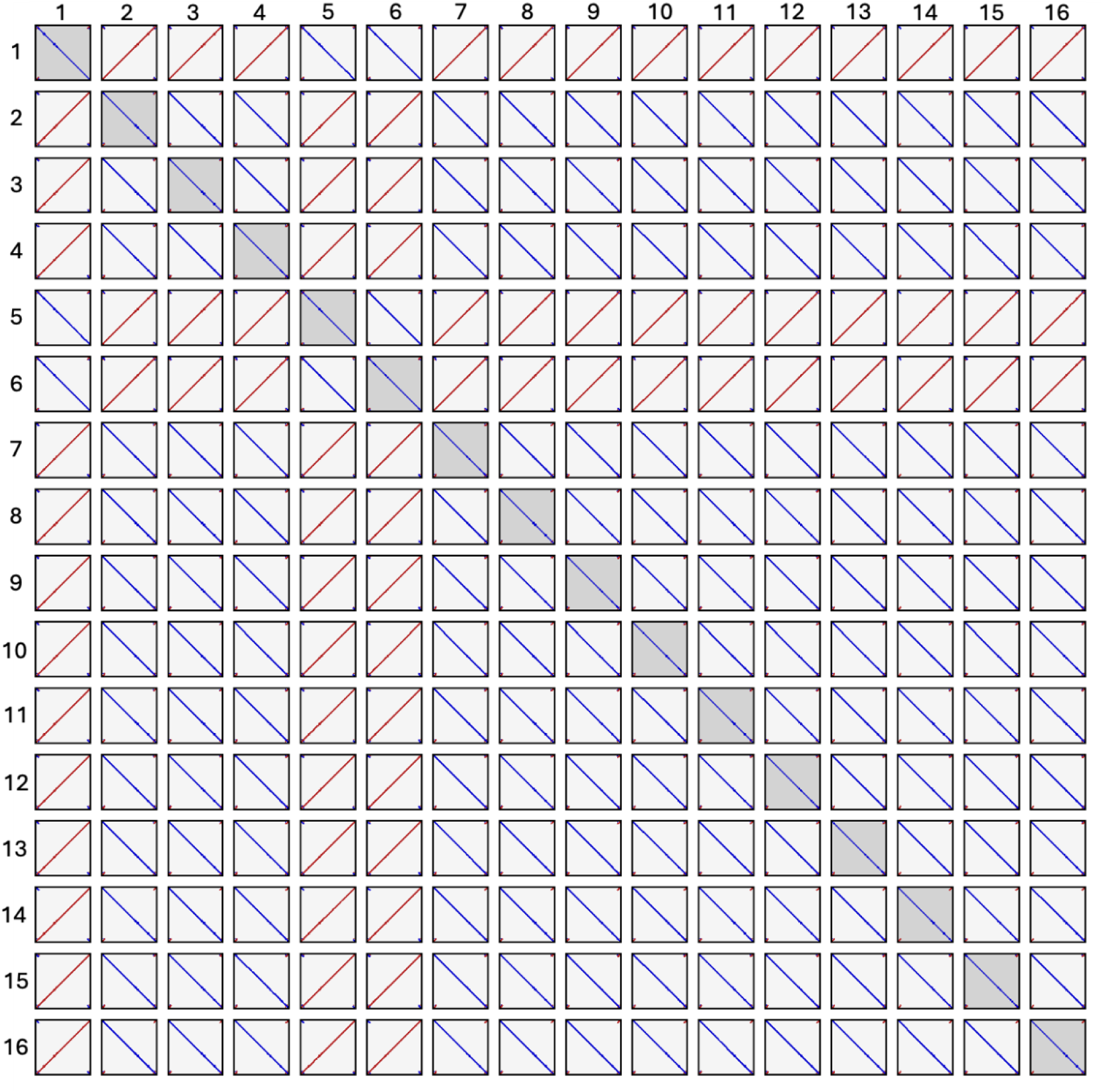
Dotplot Analysis of 16 Assembled Contigs. Squares are a visual comparison of all pairwise combinations of the 16 assembled contigs. Diagonal blue lines indicate the sequences have the same start and end sites, are highly similar, and do not contain any large gaps or rearrangements. The diagonal red lines indicate the sequences are the reverse of one another, are highly similar, and do not contain large gaps or rearrangements.

A multiple sequence alignment was generated from the remaining 16 clustered and reconciled contigs with the MUSCLE algorithm (21). Long reads were partitioned to each assembly to determine the single best alignment for each read. Then, a linear consensus sequence was generated based on the best read alignments. Long-read correction with the Nanopore tool Medaka v1.11.1 was used to polish the consensus sequence (22,23). Short Illumina read pairs were quality filtered using Fastp v0.23.4. Then, two iterations of polypolish v0.5.0 were used to correct genome errors using the filtered Illumina reads and generate the final genome assembly (24,25). Alignment of Illumina reads to the assembled genome produced a mean read depth of 21,688.3x, corrected 54 positions in the genome, and identified a 104 bp region at the C-terminus that had 0% coverage. This 104 bp region was removed from the final assembly, and likely arose from erroneous read extension. The final assembly resulted in a genome length of 165,454 bp.

### Primer Design, PCR Amplification, and Analysis of Random Genomic Regions

The CpV-BQ1 assembly was verified by PCR amplification and sequencing of five regions, approximately 9,000 bp in length, across the genome (Figure 3). To design primers, the CpV-BQ1 genome was uploaded to PrimalScheme and an amplicon size of 9,000 bp was selected (26). PrimalScheme generated 19 potential primer pairs that could be used across the entire genome, of which five were selected to perform random spot checks across the genome (Supplementary Table 1). An additional four primers were designed to sequence across the genome ends in order to investigate the genome topology (Figure 3). PCR amplification was performed with 2x GB-AMP PaCeR HP Master Mix (GeneBio Systems, PCR-002-01), 0.4 µM of forward and reverse primers, and ∼1.1 ng of genomic DNA. Thermocycling conditions were: initial denaturation at 95°C for 30 seconds, 35 cycles of 95°C for 15 s, 68°C for 15 s, and 72°C for 10 mins, then a final extension at 72°C for 6.5 min and samples were held at 12°C. PCR products were analyzed by gel electrophoresis on a 1% agarose gel and concentration was evaluated with a Qubit 4.0 fluorometer. PCR fragments were cleaned up with AMPure XP beads (Beckman-Coulter, A63881) to remove primers and small DNA fragments. Recovered products were prepared using the Nanopore Rapid Barcoding kit V14 (Nanopore, SQK-RBK114.24). The prepared library was loaded onto a MinION R10.4.1 flowcell (Nanopore, FLO-MIN114) and sequenced on a Nanopore MinION. Reads were analyzed using the Trycycler pipeline as previously described with a few exceptions. Reads were filtered once using filtlong, twelve read subsets were generated using a genome size of 9,000 bp, and only one round of polishing was performed with polypolish. Assembled contigs (Supplementary Table 2) were aligned to the CpV-BQ1 genome using the MUSCLE algorithm and visualized with Seaview (21,27).

**Figure 3.**
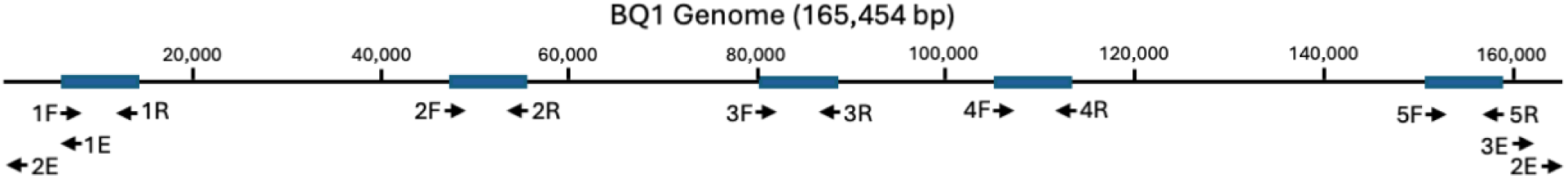
Diagram of CpV-BQ1 Genome Primer Binding Sites. The complete 165,454 bp CpV-BQ1 genome with the primer binding sites used for genome validation. Primer binding sites are represented by arrows specific to the direction of amplification. Primer pairs 1F&1R, 2F&2R, 3F&3R, 4F&4R, and 5F&5R used to amplify 9,000 bp regions in the genome indicated by blue rectangles. Primer pairs 1E&2E, 1E&3E, 2E&2E, and 2E&3E were used to PCR amplify across the genome ends (Supplementary Table 1).

### Open Reading Frame (ORF) Prediction

Open reading frames (ORFs) were predicted using a combination of four prediction tools: GeneMarkS, Glimmer3, Prodigal, and FragGeneScan (28–31). Any ORF predicted by two or more tools was considered a putative gene encoding ORF (32–34). When differing gene start sites were identified, the positions identified with GeneMarkS were used first, followed by Prodigal, then FragGeneScan (33). A total of 193 ORFs were predicted, of which GeneMarkS, Prodigal, FragGeneScan, and Glimmer3 predicted 98.45%, 95.85%, 88.08%, and 97.4% of these ORFs, respectively (Supplementary Table 3).

### Gene annotation

ORFs were assessed for functional predictions using multiple approaches, including database searches, InterPro analysis, and HMM classification. A BLASTp search against the non-redundant protein sequence database was performed twice, once including all organisms and a second search against only viruses (35). Additionally, a search was performed against the IMG virus protein BLAST database (36). InterPro was used to predict protein functionality using protein domain and family predictions (37). InterPro has multiple tools integrated into its online platform including Panther, NCBIfam, CDD, Cath-Gene3D, SUPERFAMILY, ProSiteProfiles, ProSitePatterns, PRINTS, SMART, Pfam, HHMAP, and FunFam to classify protein families, domains, superfamilies, and amino acid sites. InterPro also identifies signal peptides, transmembrane helices, coils, and disordered regions using TMHMM, Coils, MobliteDB, and Phobius. HMM profile analysis against the PhROGs, efam-xc, RVDB, and VOGDB were performed to classify ORFs (38–41). MMSeqs was used to perform HMM analysis against the PhROGs database while HMMer3 was used to search all other HMM databases (42,43). tRNAscan-SE was used to predict transfer RNA (tRNA) sequences in the assembled CpV-BQ1 genome (44).

### Taxonomic classification

Three genes common to all NCLDVs were used to taxonomically classify the CpV-BQ1 genome, DNA polymerase type-B, viral A32-like packaging ATPase, and viral late transcription factor 3 (VLTF-3). Protein sequences and accessions for each gene from various NCLDVs were downloaded from NCBI (Supplementary Table 4). Multiple sequence alignments (MSAs) were generated using the MUSCLE algorithms for each protein using Seaview v5.0.4 (21,27). MSAs were used to generate phylogenetic trees in Seaview using the phylogenetic maximum likelihood method with the LG model, aLRT (SH-like) branch support, NNI tree searching operation, and a BioNJ starting tree (27,45).

## Results and Discussion

### Assembly of the CpV-BQ1 Genome

A hybrid assembly approach with long Nanopore and short Illumina reads was used to recover the CpV-BQ1 genome. Long read assemblies can produce larger contigs and resolve repetitive genome regions with higher accuracy than short Illumina read assemblies. However, Illumina reads have higher sequence accuracy than long reads and can correct for errors introduced through long read sequencing. Thus, using a combination of long and short reads for assembly produces a genome with higher accuracy and instills greater confidence in the resulting assembly (16,46). Hybrid assembly of the CpV-BQ1 genome was performed using the TryCycler pipeline (16). TryCycler is a robust tool that uses a unique approach to produce assemblies with a high degree of confidence. Most long-read assembly tools work fairly well, however, they are not perfect and can introduce large- and small-scale errors into an assembled genome which often go undetected (16). To eliminate these issues, TryCycler uses multiple separate long-read assemblies, generated by a variety of assembly tools, as its input and produces a consensus sequence. By using a variety of assembly tools which utilize different approaches/algorithms, bias and/or errors introduced by any one tool can be detected and eliminated (16).

The TryCycler pipeline begins by randomly dividing long reads into discrete subsets. Read subsets are then assembled into contigs using differing assembly tools. With deep enough sequencing, each subset should contain reads that cover the complete genome, thus highly similar contigs should be produced from the assembly of each read subset. Next, a phylogenetic analysis is performed which is used to generate clusters of closely related contigs. Highly accurate assemblies form clear phylogenetic clusters and exclude assemblies which contain spurious, incomplete, or misassembled contigs (16). If clear clusters are not formed the assemblies are considered inconsistent and unreliable. High quality clustered contigs are then reconciled to ensure sequence similarity and an attempt to circularize assemblies is made. A multiple sequence alignment (MSA) is generated from reconciled contigs then reads are partitioned to each contig to determine the single best alignment for each read. Using the MSA and best read alignments, TryCycler generates a consensus sequence resolving differences between assemblies and producing the most likely genome assembly. Finally, high quality short Illumina reads are aligned to the assembled genome and used to correct long-read sequencing errors, producing a highly accurate and trust-worthy final genome assembly (16).

The cleaned and filtered long reads were divided into 24 subsets. Then, six subsets were assembled with Flye, Minipolish, Raven, and Canu, respectively (17–20). A total of 26 contigs were assembled of which 23 formed a single phylogenetic cluster (Figure 1). After reconciling these assemblies, 16 contigs showed high sequence similarity. This included five Flye, and Minipolish assemblies, four Raven assemblies, and two Canu assemblies (Figure 1). The similarity between the 16 independently generated contigs provided confidence that the assemblies were of very high quality and representative of the true CpV-BQ1 genome. These 16 contigs were used to generate an MSA, reads were partitioned, then a consensus genome was generated and polished with Illumina reads producing a 165,454 bp genome. Additionally, attempts to circularize the genome during reconciling were unsuccessful. Analysis of these contigs using a dotplot shows the same discrete start and end point in all 16 contigs which indicates the genome has a linear topology (Figure 2). If genomes had a circular topology, we would expect the start and stop sites to vary amongst the 16 assembled genomes, causing the lines within each dotplot to start along the axis edge instead of the corner. Additionally, if some of our assemblies contained gaps or large rearrangements the plots would contain lines with gaps or a discontinuous arrangement. Our dotplot indicates these 16 assemblies are not circular and do not contain gaps or large rearrangements (Figure 2).

To validate the assembled sequence PCR amplifications, Nanopore sequencing and TryCycler assembly were performed to recover and verify the sequence of five 9,000 bp regions within the genome (Figure 3). Additionally, to verify the genomes linear topology, four PCR amplifications across the genome ends were attempted (Figure 3).

The five ∼9,000 bp regions were easily PCR amplified, sequenced, and assembled with TryCycler (Supplementary Table 2). Alignment of the assembled contigs to the CpV-BQ1 genome showed almost 100% sequence identity to their respective sites. The very ends of assembled PCR regions # two and # five, contained one and 17 bases that differ from the CpV-BQ1 genome, respectively, which can be attributed to low read coverage in these areas. Overall, the similarity of PCR amplified sequences to the assembled genome provides further confidence in the assembly accuracy. Attempts to amplify genome ends to test for circularization were unsuccessful. Of these four attempted PCR amplifications (Figure 3), nanopore reads were obtained from three reactions. However, attempts at read assembly with TryCycler resulted in clusters with many spurious, incomplete, and misassembled contigs. Thus, the inability to amplify sequences which span the genome ends provides additional confirmation that the CpV-BQ1 genome has a linear topology.

### Characterization and Description of the CpV-BQ1 Genome

Assembly of the CpV-BQ1 genome sequence produced a linear genome of 165,454 bp with a GC content of 32.32%. Three tRNA sequences and 193 coding sequences (CDSs) were identified in the genome which ranged from 44 to 1693 amino acids (aa) in length, with an average size of 260 aa (Figure 4, Supplementary Table 3). Coding sequences were found in slightly higher prevalence on the positive strand than the negative strand, occurring at a rate of 58.5% and 41.5%, respectively. Of the 193 identified CDSs, 92 (47.67%) were assigned putative functions based on homology to known genes and protein domains, while the remaining 101 (52.33%) could not be assigned functions and were designated as hypothetical proteins (Supplementary Table 3). Functionally annotated CDSs were placed into eight general functional groups (Figure 4, Table 1). A total of 19 CDSs were involved in DNA replication, recombination, and repair; eight in nucleotide metabolism and DNA packaging; 10 in transcription; three in sugar manipulation; four in DNA methylation; 15 in protein and lipid binding, synthesis, and modification; 12 in virion capsid and associated proteins, and 21 encoded miscellaneous functions (Table 1).

**Figure 4.**
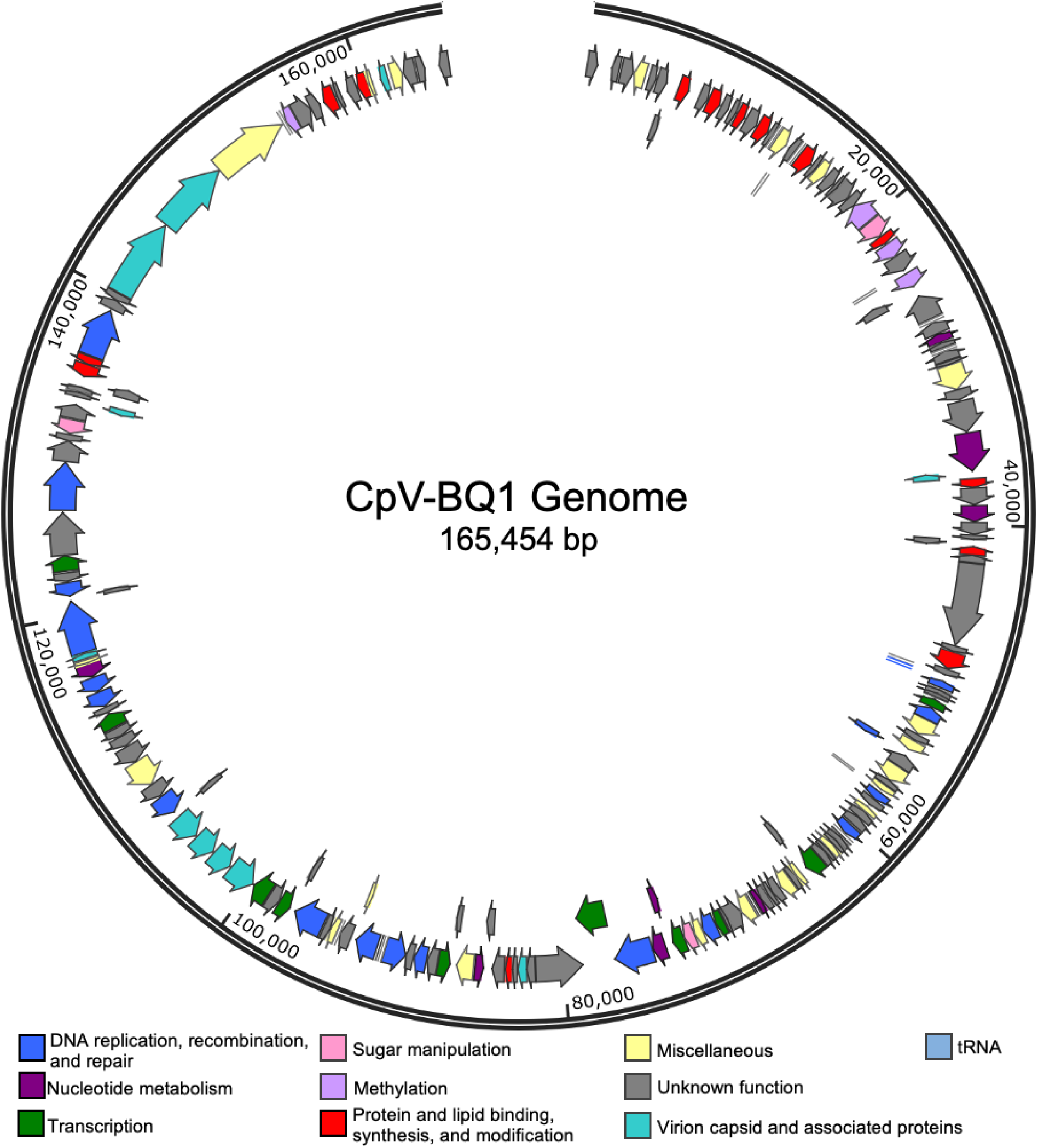
ORF Predictions in the CpV-BQ1 Genome. ORFs represented by arrows are aligned to the 165,454 bp CpV-BQ1 genome oriented in their coding direction. Arrows are coloured based on their hypothesized functional group. Blue arrows for DNA replication, recombination, and repair; dark purple for nucleotide metabolism and DNA packaging; green for transcription; pink for sugar manipulation; lavender for methylation; red for protein and lipid binding, synthesis, and modification; aqua for virion capsid and associated proteins; yellow for miscellaneous proteins; light blue for tRNA; and grey for all proteins with unknown functions. The genome is presented as circular for presentation purposes only. Nucleotide positions are denoted at every 20,000 bps.

**Table 1.**
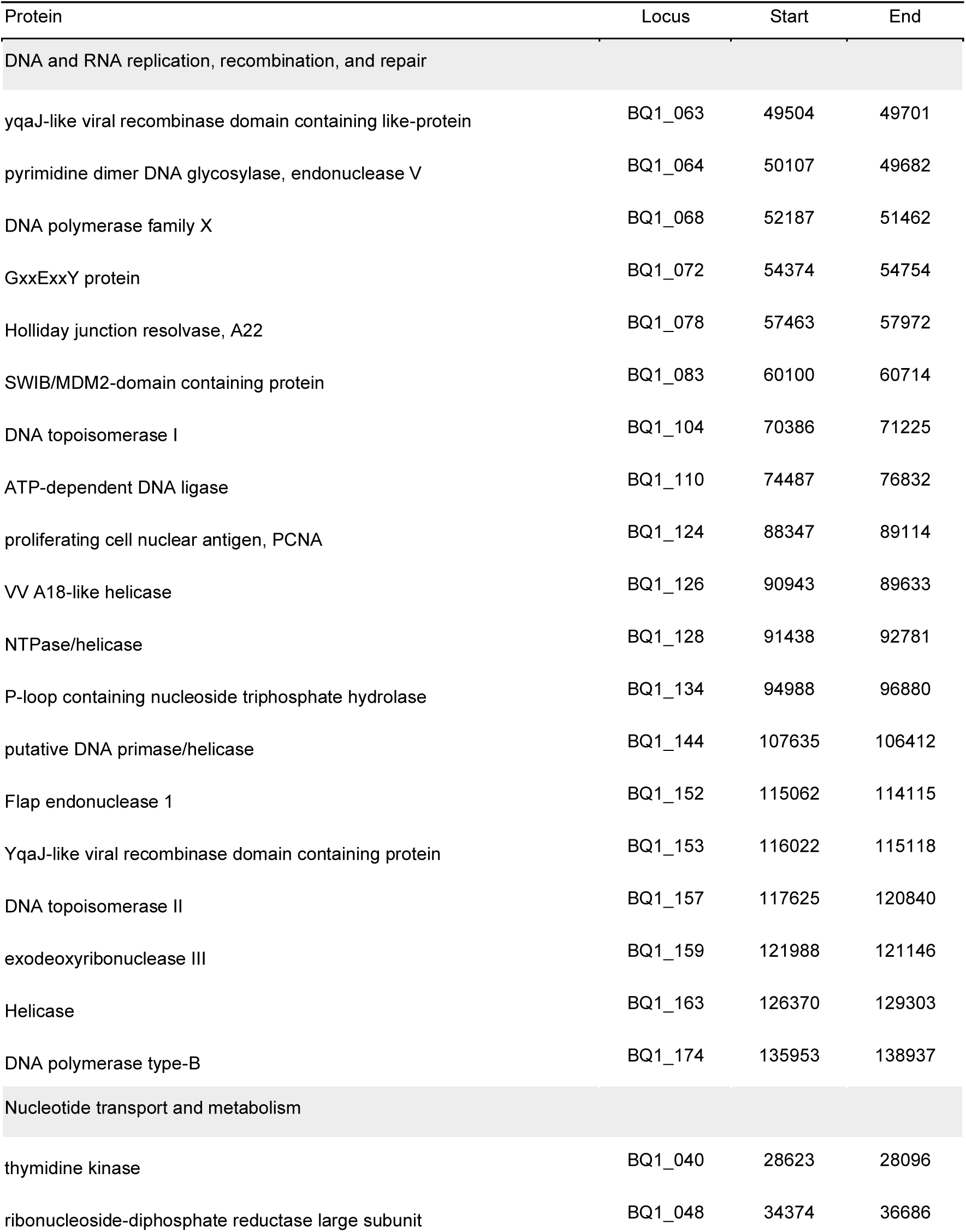
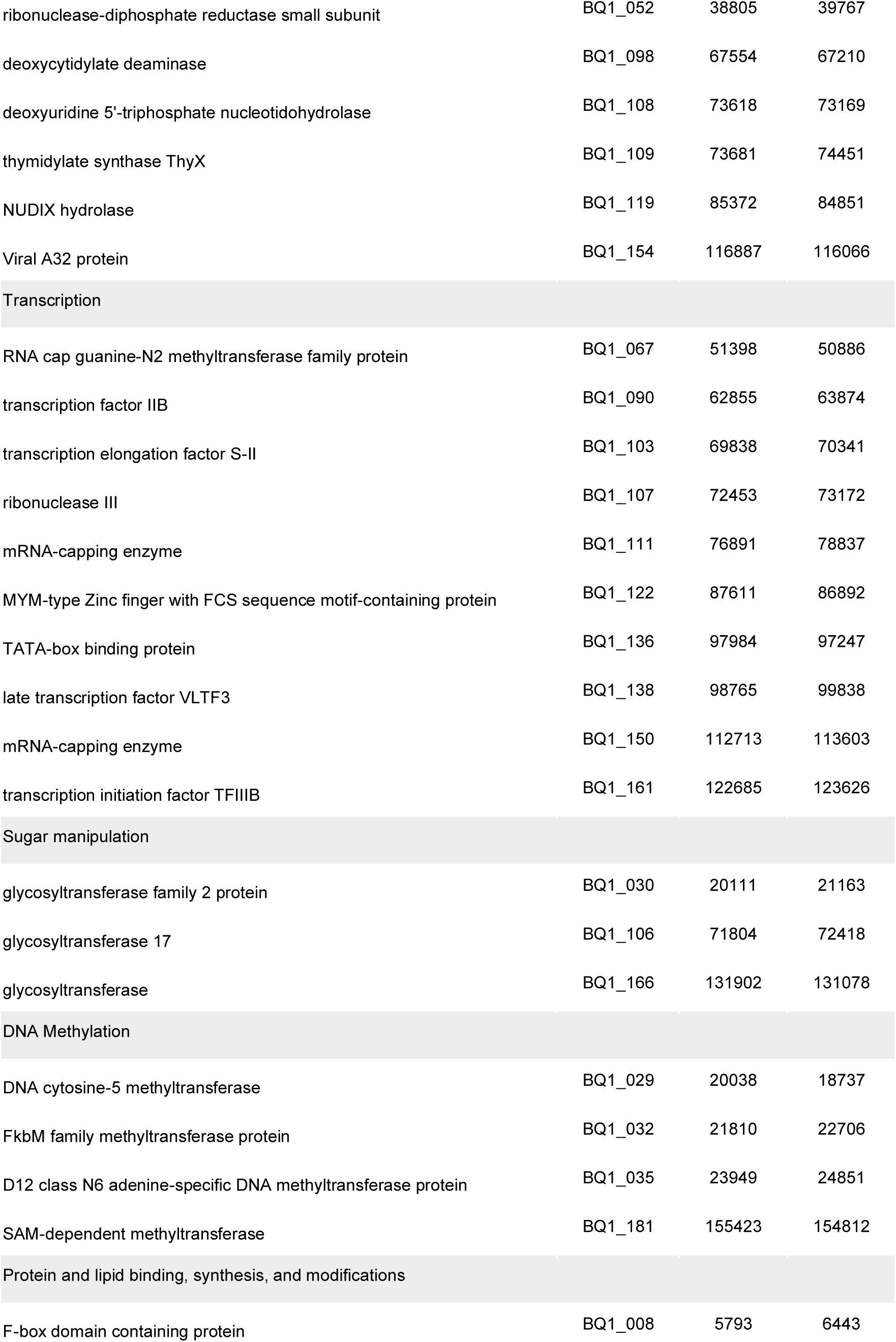
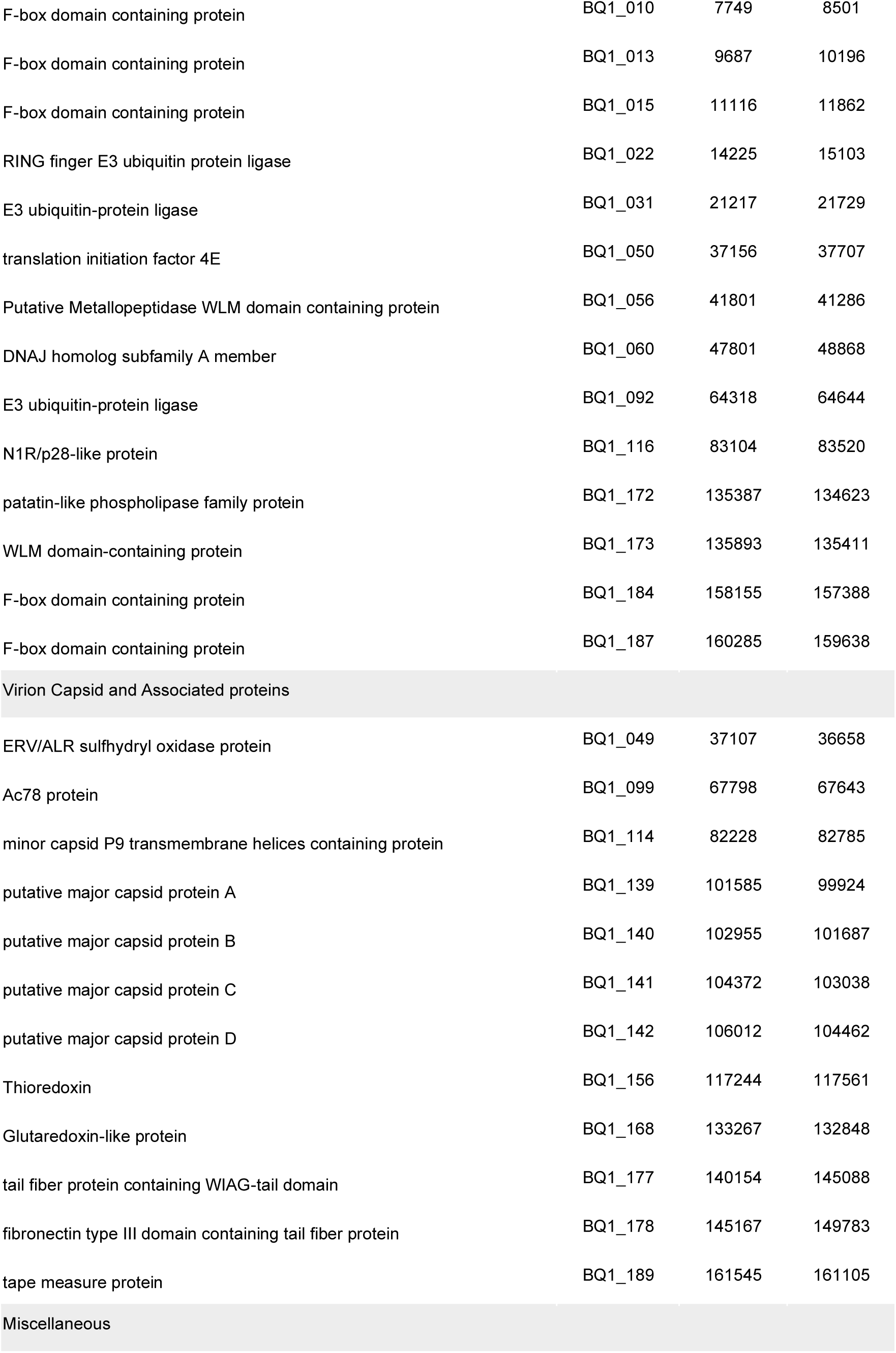
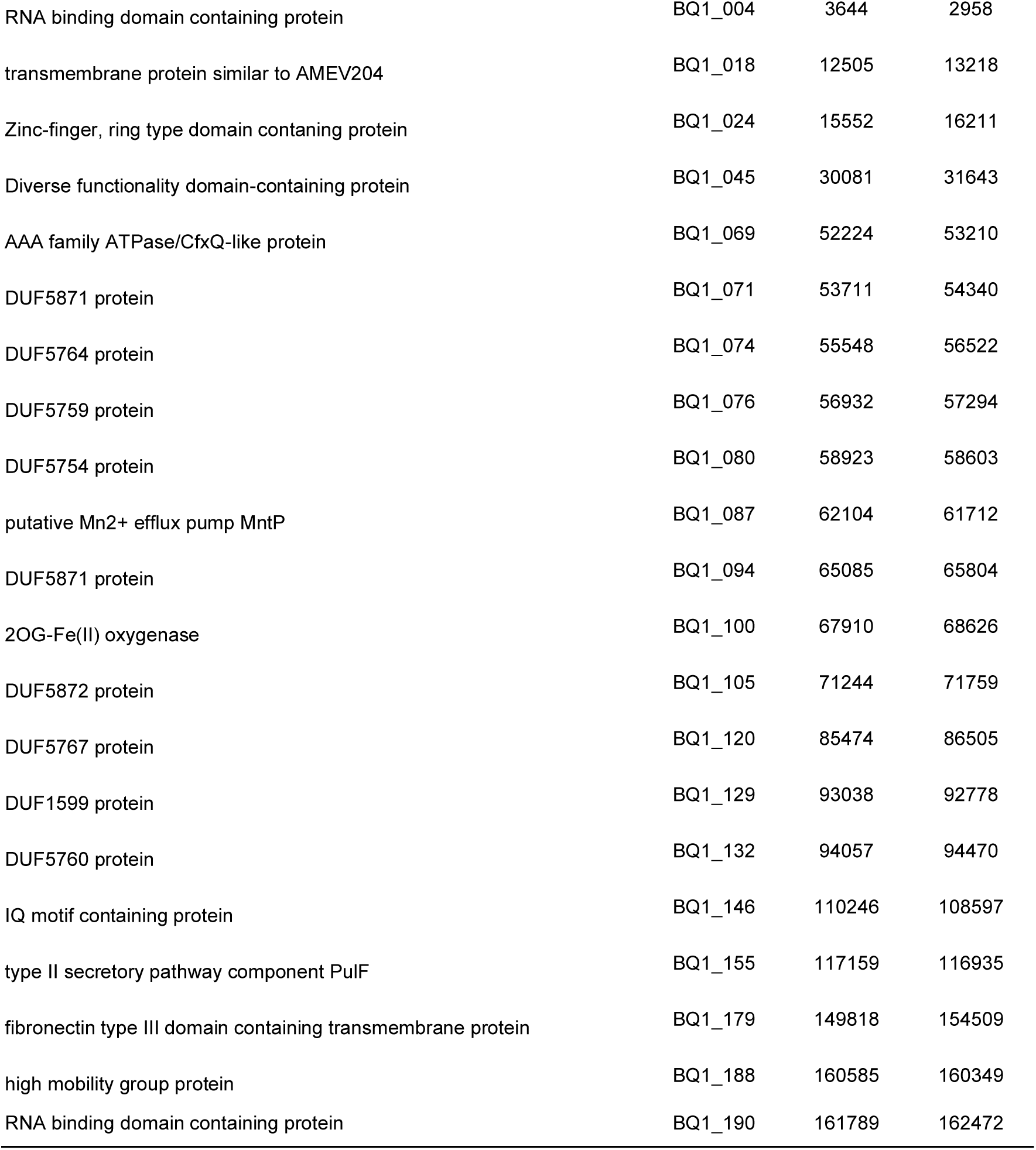
List of all Characterised Genes in the CpV-BQ1 Genome. All genes in the CpV-BQ1 genome with an identifiable function or domain architecture. Gene name, locus, as well as start and end position within the CpV-BQ1 genome are provided. These protein encoding genes are divided into functional groups including, DNA replication, recombination, and repair; nucleotide metabolism and DNA packaging; transcription; sugar manipulation; DNA methylation; protein and lipid binding, synthesis, and modifications; virion capsid and associated proteins; and miscellaneous.

| Protein | Locus | Start | End |
| --- | --- | --- | --- |
| DNA and RNA replication, recombination, and repair |  |  |  |
| yqaj-like viral recombinase domain containing like-protein | BQ1_063 | 49504 | 49701 |
| pyrimidine dimer DNA glycosylase, endonuclease V | BQ1_064 | 50107 | 49682 |
| DNA polymerase family X | BQ1_068 | 52187 | 51462 |
| GxxExxY protein | BQ1_072 | 54374 | 54754 |
| Holliday junction resolvase, A22 | BQ1_078 | 57463 | 57972 |
| SWIB/MDM2-domain containing protein | BQ1_083 | 60100 | 60714 |
| DNA topoisomerase I | BQ1_104 | 70386 | 71225 |
| ATP-dependent DNA ligase | BQ1_110 | 74487 | 76832 |
| proliferating cell nuclear antigen, PCNA | BQ1_124 | 88347 | 89114 |
| VV A18-like helicase | BQ1_126 | 90943 | 89633 |
| NTPase/helicase | BQ1_128 | 91438 | 92781 |
| P-loop containing nucleoside triphosphate hydrolase | BQ1_134 | 94988 | 96880 |
| putative DNA primase/helicase | BQ1_144 | 107635 | 106412 |
| Flap endonuclease 1 | BQ1_152 | 115062 | 114115 |
| Yqaj-like viral recombinase domain containing protein | BQ1_153 | 116022 | 115118 |
| DNA topoisomerase II | BQ1_157 | 117625 | 120840 |
| exodeoxyribonuclease III | BQ1_159 | 121988 | 121146 |
| Helicase | BQ1_163 | 126370 | 129303 |
| DNA polymerase type-B | BQ1_174 | 135953 | 138937 |
| Nucleotide transport and metabolism |  |  |  |
| thymidine kinase | BQ1_040 | 28623 | 28096 |
| ribonucleoside-diphosphate reductase large subunit | BQ1_048 | 34374 | 36686 |
| ribonuclease-diphosphate reductase small subunit | BQ1_052 | 38805 | 39767 |
| deoxycytidylate deaminase | BQ1_098 | 67554 | 67210 |
| deoxyuridine 5'-triphosphate nucleotidohydrolase | BQ1_108 | 73618 | 73169 |
| thymidylate synthase ThyX | BQ1_109 | 73681 | 74451 |
| NUDIX hydrolase | BQ1_119 | 85372 | 84851 |
| Viral A32 protein | BQ1_154 | 116887 | 116066 |
| Transcription |  |  |  |
| RNA cap guanine-N2 methyltransferase family protein | BQ1_067 | 51398 | 50886 |
| transcription factor IIB | BQ1_090 | 62855 | 63874 |
| transcription elongation factor S-II | BQ1_103 | 69838 | 70341 |
| ribonuclease III | BQ1_107 | 72453 | 73172 |
| mRNA-capping enzyme | BQ1_111 | 76891 | 78837 |
| MYM-type Zinc finger with FCS sequence motif-containing protein | BQ1_122 | 87611 | 86892 |
| TATA-box binding protein | BQ1_136 | 97984 | 97247 |
| late transcription factor VLTF3 | BQ1_138 | 98765 | 99838 |
| mRNA-capping enzyme | BQ1_150 | 112713 | 113603 |
| transcription initiation factor TFIIIB | BQ1_161 | 122685 | 123626 |
| Sugar manipulation |  |  |  |
| glycosyltransferase family 2 protein | BQ1_030 | 20111 | 21163 |
| glycosyltransferase 17 | BQ1_106 | 71804 | 72418 |
| glycosyltransferase | BQ1_166 | 131902 | 131078 |
| DNA Methylation |  |  |  |
| DNA cytosine-5 methyltransferase | BQ1_029 | 20038 | 18737 |
| FkbM family methyltransferase protein | BQ1_032 | 21810 | 22706 |
| D12 class N6 adenine-specific DNA methyltransferase protein | BQ1_035 | 23949 | 24851 |
| SAM-dependent methyltransferase | BQ1_181 | 155423 | 154812 |
| Protein and lipid binding, synthesis, and modifications |  |  |  |
| F-box domain containing protein | BQ1_008 | 5793 | 6443 |
| F-box domain containing protein | BQ1_010 | 7749 | 8501 |
| F-box domain containing protein | BQ1_013 | 9687 | 10196 |
| F-box domain containing protein | BQ1_015 | 11116 | 11862 |
| RING finger E3 ubiquitin protein ligase | BQ1_022 | 14225 | 15103 |
| E3 ubiquitin-protein ligase | BQ1_031 | 21217 | 21729 |
| translation initiation factor 4E | BQ1_050 | 37156 | 37707 |
| Putative Metallopeptidase WLM domain containing protein | BQ1_056 | 41801 | 41286 |
| DNAJ homolog subfamily A member | BQ1_060 | 47801 | 48868 |
| E3 ubiquitin-protein ligase | BQ1_092 | 64318 | 64644 |
| N1R/p28-like protein | BQ1_116 | 83104 | 83520 |
| patatin-like phospholipase family protein | BQ1_172 | 135387 | 134623 |
| WLM domain-containing protein | BQ1_173 | 135893 | 135411 |
| F-box domain containing protein | BQ1_184 | 158155 | 157388 |
| F-box domain containing protein | BQ1_187 | 160285 | 159638 |
| Virion Capsid and Associated proteins |  |  |  |
| ERV/ALR sulfhydryl oxidase protein | BQ1_049 | 37107 | 36658 |
| Ac78 protein | BQ1_099 | 67798 | 67643 |
| minor capsid P9 transmembrane helices containing protein | BQ1_114 | 82228 | 82785 |
| putative major capsid protein A | BQ1_139 | 101585 | 99924 |
| putative major capsid protein B | BQ1_140 | 102955 | 101687 |
| putative major capsid protein C | BQ1_141 | 104372 | 103038 |
| putative major capsid protein D | BQ1_142 | 106012 | 104462 |
| Thioredoxin | BQ1_156 | 117244 | 117561 |
| Glutaredoxin-like protein | BQ1_168 | 133267 | 132848 |
| tail fiber protein containing WIAG-tail domain | BQ1_177 | 140154 | 145088 |
| fibronectin type III domain containing tail fiber protein | BQ1_178 | 145167 | 149783 |
| tape measure protein | BQ1_189 | 161545 | 161105 |
| Miscellaneous |  |  |  |
| RNA binding domain containing protein | BQ1_004 | 3644 | 2958 |
| transmembrane protein similar to AMEV204 | BQ1_018 | 12505 | 13218 |
| Zinc-finger, ring type domain containing protein | BQ1_024 | 15552 | 16211 |
| Diverse functionality domain-containing protein | BQ1_045 | 30081 | 31643 |
| AAA family ATPase/CfxQ-like protein | BQ1_069 | 52224 | 53210 |
| DUF5871 protein | BQ1_071 | 53711 | 54340 |
| DUF5764 protein | BQ1_074 | 55548 | 56522 |
| DUF5759 protein | BQ1_076 | 56932 | 57294 |
| DUF5754 protein | BQ1_080 | 58923 | 58603 |
| putative Mn<sup>2+</sup> efflux pump MntP | BQ1_087 | 62104 | 61712 |
| DUF5871 protein | BQ1_094 | 65085 | 65804 |
| 2OG-Fe(II) oxygenase | BQ1_100 | 67910 | 68626 |
| DUF5872 protein | BQ1_105 | 71244 | 71759 |
| DUF5767 protein | BQ1_120 | 85474 | 86505 |
| DUF1599 protein | BQ1_129 | 93038 | 92778 |
| DUF5760 protein | BQ1_132 | 94057 | 94470 |
| IQ motif containing protein | BQ1_146 | 110246 | 108597 |
| type II secretory pathway component PulF | BQ1_155 | 117159 | 116935 |
| fibronectin type III domain containing transmembrane protein | BQ1_179 | 149818 | 154509 |
| high mobility group protein | BQ1_188 | 160585 | 160349 |
| RNA binding domain containing protein | BQ1_190 | 161789 | 162472 |

### Taxonomy

Viruses within the *Megaviricetes* (previously and informally known as NCLDVs) share a few common characteristics, including the presence of a dsDNA genome typically over 100 kb, which encodes a handful of universal NCLDV genes (3). These genes include the major capsid protein (MCP), a DNA polymerase B (polB), the viral A32-like packing ATPase, and the viral late transcription factor 3 (VLTF-3) (3). Additionally, although the D5-helicase is considered a universal NCLDV gene, it is not found in viruses in the *Phycodnaviridae* family, nor in the CpV-BQ1 genome (3). Of the universal genes, the MCPs share little sequence identity, and thus cannot be used reliably for phylogenetic analysis. Phylogenetic analysis of the CpV-BQ1 genome was therefore performed using polB, viral A32-like packing ATPase, and VLTF-3 (Figure 5). In all three phylogenetic trees, the CpV-BQ1 groups closely with viruses in the *Phycodnaviridae* family (Figure 5) such as the Heterosigma akashiwo virus 01, a Phycodnaviridae virus and the single member of the genus *Raphidovirus*. Within the DNA polymerase B phylogenetic tree CpV-BQ1 groups closest to two viruses within the genus *Prymnesiovirus* (47), Chrysochromulina brevifilum virus PW1 (CbV-PW1) (48) and Phaeocystis globosa virus 08T (PgV-08T) (49). Molecular analysis of the A32 ATPase and VLTF-3 genes in CbV-PW1 and PgV-08T has not been performed, thus we could not include them in our phylogenetic analysis. Nonetheless, based on the similarity of the polB gene and the fact these three viruses all have prymnesiophyte hosts (47) suggests the most appropriate classification for CpV-BQ1 is within the *Prymnesiovirus* genus. Thus, our phylogenetic analysis supports the inclusion of CpV-BQ1 in the NCLDV *Phycodnaviridae* family and suggests it can most appropriately be placed within the *Prymnesiovirus* genus.

**Figure 5.**
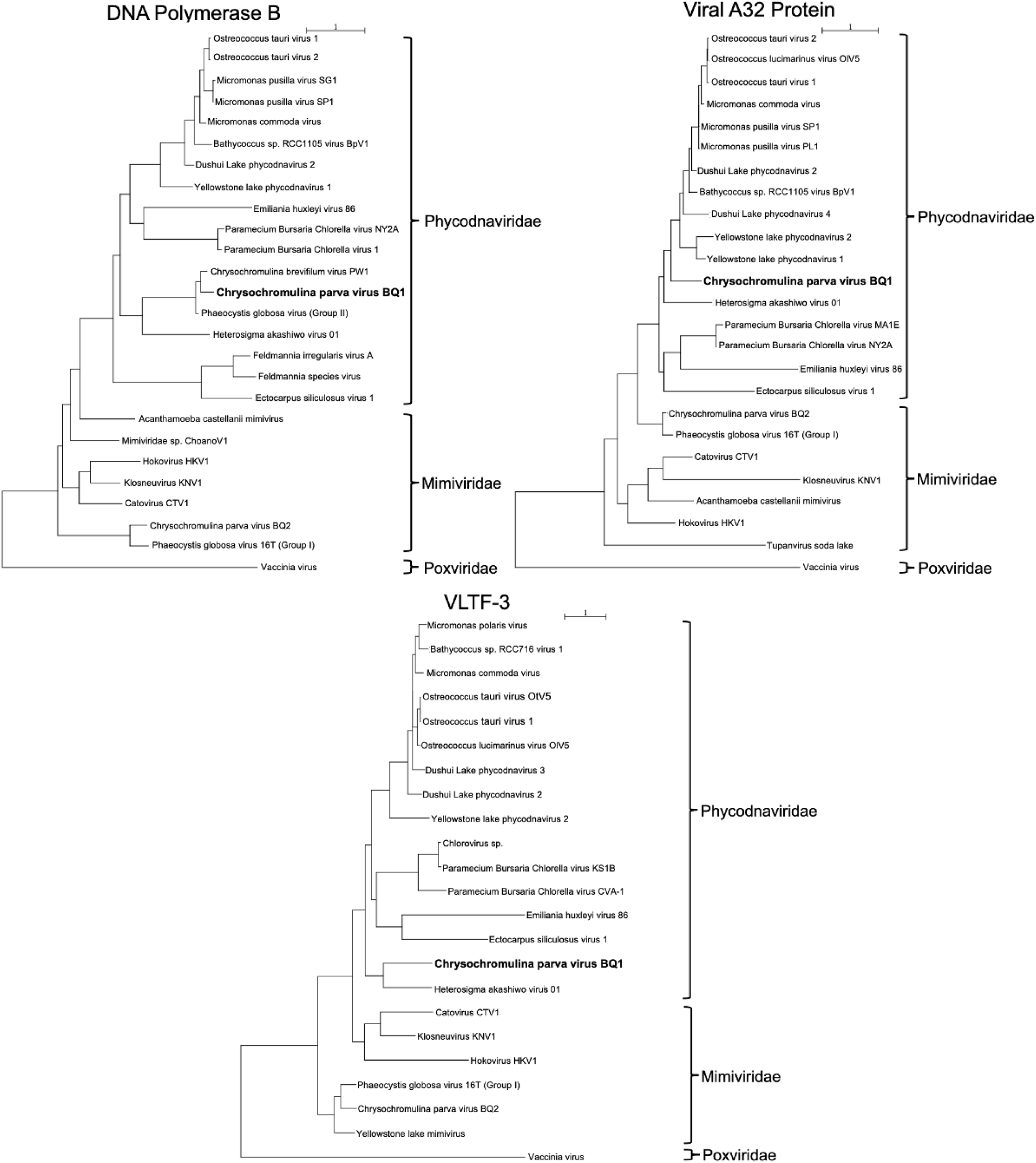
Phylogenetic analysis of three CpV-BQ1 genes universally found in NCLDV genomes. Phylogenetic trees generated to determine the taxonomy of the CpV-BQ1 virus using the Phylogenetic maximum likelihood (PhyML) algorithm in Seaview v5.0. Trees were generated using the protein sequences for the universal NCLDV genes polB, viral A32 protein, and VLTF-3 (Supplementary Table 4). Protein sequences are from viruses belonging to either the *Phycodnaviridae*, *Mimiviridae*, or *Poxviridae* family.

### DNA and RNA replication, recombination, and repair enzymes

NCLDVs encode a suite of genes which enable the replication of their own genome with little reliance on their host’s genome replication machinery (50,51). A total of 19 proteins involved in DNA/RNA replication, recombination, and repair were identified in the CpV-BQ1 genome. This consort of enzymes enables CpV-BQ1 to facilitate much of its own DNA replication processes. These replication associated genes include a DNA polymerase type-B (polB), a DNA polymerase family X protein, an A22 holliday junction resolvase, a proliferating cell nuclear antigen sliding clamp (PCNA), topoisomerase I and II, four nuclease proteins, four helicase proteins, two YqaJ-like viral recombinases, an ATP-dependant DNA-ligase, a P-loop nucleoside triphosphate hydrolase, and a SWIIB/MDM2-domain protein (50,51). Some NCLDVs, such as Poxviruses and Mimiviruses, replicate and package their genomes within the cytoplasm of their host cell in subcellular compartments surrounded by endoplasmic reticulum membranes, which are referred to as virus factories (50,52). However, other NCLDVs, such as Phycodnaviridae viruses, replicate their genome within the nucleus of their host, then package virions in the cytoplasmic virus factories (52,53). To explore the location of CpV-BQ1 genome replication, proteins involved in DNA replication were analyzed for the presence of nuclear localization signals (NLS) using DeepLoc 2.1 (54). Indeed, NLS signals were identified in 12 of the 19 predicted DNA replication, recombination, and repair enzymes, including the DNA-dependent DNA replication polymerase, polB. This suggests that at least some part of the DNA replication process occurs within the host cell nucleus.

### Transcription

Most NCLDVs encode a DNA-dependent RNA polymerase which allows them to transcribe their viral mRNA within the cytoplasm (55). However, some members of the *Phycodnaviridae* family, such as *Chloroviruses* and *Prasinoviruses,* do not encode an RNA polymerase gene. Instead they rely on the host’s RNA polymerase and enter the nucleus to initiate viral gene transcription (56). Similarly, the CpV-BQ1 genome does not encode an recognizable RNA polymerase and therefore its lytic cycle most likely involves a nuclear localization step necessary for gene transcription.

To gain control of the host RNA polymerase II (RNA pol), CpV-BQ1 encodes several eukaryotic-like transcription factors (TFs) which likely interact with the host’s RNA pol and alter its activity. CpV-BQ1 encodes two transcription factor IIB (TFIIB) proteins and a TATA box binding protein (TBP), which are similar to general eukaryotic TFs (57). During eukaryotic transcription initiation, RNA pol is recruited to the DNA promoter by a complex of general TFs including TFIIB and TBP. With virus encoded TFs, viruses can redirect the RNA pol to initiate the transcription of viral genes (57). CpV-BQ1 also encodes two specific TFs which activate viral genes expressed in the late transcription phase, including virus late transcription factor 3 (VLT-3) and an MYM-type zinc-finger FCS motif containing protein (58–60).

A handful of other genes important for transcription are encoded by the CpV-BQ1 genome. Transcription elongation factor S-II (TFIIS) stimulates the cleavage activity of RNA pol when the transcription complex becomes stalled, allowing transcription to be restarted at the newly created 3’ prime end (61). Two mRNA-capping enzymes are encoded which ensure the protection and efficient transcription of mRNA. A protein Blast analysis of these two mRNA-capping enzymes show they may differ in their evolutionary origin and possibly function. The first is encoded at locus BQ1_111 and is similar to bacterial and eukaryotic mRNA capping enzymes, while the second locus is similar to archaeal and viral mRNA capping enzymes. Moreover, an RNA cap guanine-N2 methyltransferase family protein likely methylates the 5’ prime cap on mRNA, which is suspected to increase the synthesis of viral transcripts (62). Lastly, the enzyme ribonuclease III (RNase III) is encoded by the CpV-BQ1 genome. RNase III proteins are involved in the processing and maturation of RNA species, including tRNA, rRNA, and mRNA, as well as the degradation of mRNAs. The specific role of RNase III encoded by NCLDVs is not well understood, however it is suspected that it is involved in RNA cleavage and tRNA maturation processes (63).

During the nuclear infection stage, chloroviruses such as Paramecium bursaria chlorella virus (PBCV-1) not only redirect host transcriptional machinery, but they also repress the expression of host genes (64). It is likely that CpV-BQ1 uses mechanisms similar to PBCV-1 to decrease the transcription of host genes and prioritize the expression of its own genes. In the virion, PBCV-1 packages methylation, restriction endonuclease, and chromatin-remodeling enzymes which are released upon nuclear entry of virion particles. In consort, these enzymes methylate, degrade, and remodel host DNA effectively downregulating the production of host transcripts (64). CpV-BQ1 encodes several methylase enzymes and endonucleases, some of which may target host DNA. Additionally, CpV-BQ1 encodes a SWIB/MDM2-domain containing protein; the SWIB domain, although not fully understood, can function as a transcriptional activator or as a chromatin-remodeling protein (65). Thus, circumstantial evidence suggests the CpV-BQ1 genome contains the necessary protein apparatus to downregulate host gene transcription.

### Nucleotide transport and metabolism proteins

To facilitate rapid genome replication, some NCLDVs encode DNA precursor metabolism proteins which generate pools of available deoxythymidine triphosphate (dTTP) (50). The CpV-BQ1 genome encodes several genes which are important for dTTP synthesis, including deoxycytidylate deaminase (dCD), thymidine kinase (TK), deoxyuridine 5’-triphosphate nucleotidohydrolase (dUTPase), thymidylate synthase ThyX (thyX) and ribonucleoside-diphosphate reductase (RNR) (Figure 6) (66).

**Figure 6.**
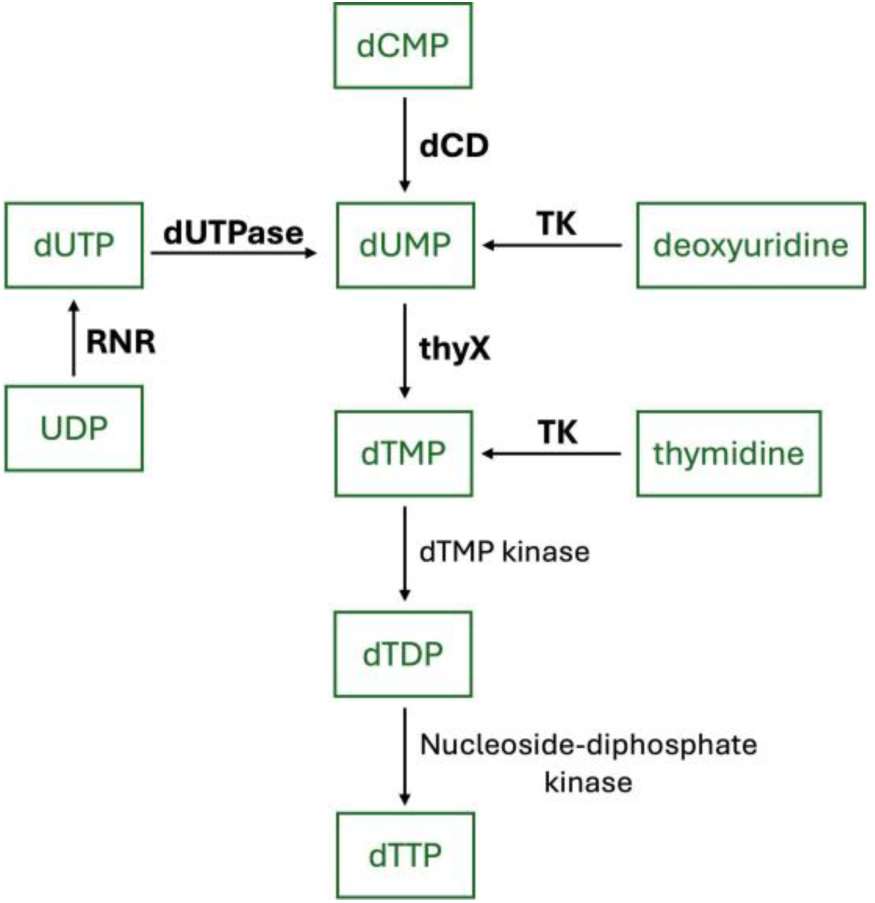
Biosynthesis of deoxythymidine triphosphate (dTTP). KEGG ontology pyrimidine metabolism pathway (https://www.kegg.jp/pathway/ec00240+3.5.4.12). Substrates are in green boxes, enzymes written in black text convert substrates following the direction of the arrow heads. Enzyme names that are bolded are encoded by the CpV-BQ1 genome, while the remaining enzymes are not encoded by the CpV-BQ1 genome.

The three enzymes dCD, TK, and dUTPase convert their respective substrates to dUMP, an essential precursor molecule for the production of dTMP by the enzyme thyX (Figure 6). Furthermore, TK can also phosphorylate available thymidine to produce dTMP (67). dTMP can then be phosphorylated by dTMP kinase then nucleoside diphosphate kinase to produce dTTP which can be incorporated into replicating viral DNA, however, the CpV-BQ1 genome does not encode either of these enzymes (Figure 6). In addition to contributing to dTTP biosynthesis, RNR can also convert CDP to dCDP with help from the protein glutaredoxin, which is encoded by the CpV-BQ1 genome. dCDP can then be phosphorylated by nucleoside-diphosphate kinase to produce dCTP which can be used for DNA synthesis. However, it has been shown that these proteins are only essential for maintaining dCTP levels when similar host enzymes have been depleted (68). Other studies have demonstrated that glutaredoxin is an important component in a redox cascade reaction that is required for virion assembly (69). Analysis of the CpV-BQ1 encoded glutaredoxin with subcellular prediction tool DeepLoc identified an endoplasmic reticulum (ER) localization signal, thus this protein is more likely to be located at ER virus factories and involved in virion assembly and packaging.

The CpV-BQ1 genome also encodes an NUDIX hydrolase. NUDIX proteins hydrolyze substrates containing a nucleoside diphosphate linked to some other moiety which increases free nucleotides available for metabolism (70). It remains unclear which substrate(s) the CpV-BQ1 encoded NUDIX hydrolase has specificity to, however it has been shown that NUDIX enzymes in NCLDVs Vaccinia Virus, African Swine Fever Virus (ASFV), and Mimivirus L375 function as mRNA decapping enzymes. This decapping process is thought to facilitate mRNA turnover and increase the availability of dNTPs (71).

### DNA Methylation

Methyltransferases (MTases) are present in all domains of life however they are not highly prevalent amongst viruses. Within viruses, MTases are most often reported in bacteriophages and some members of the *Phycodnaviridae* family (66,72). *Phycodnaviruses* often use their encoded MTases to methylate their own genome. Moreover, it has been observed that some chloroviruses encode their own restriction-modification (R-M) systems, complete with type II site specific restriction enzymes to degrade host or foreign DNA and MTases to protect their own genome from degradation (72,73). The CpV-BQ1 genome encodes five methyltransferase enzymes, including a D12 class N6 adenine-specific DNA methyltransferase protein (mA6-MTase), DNA cytosine-5 methyltransferase (mC5-MTase), FbkM methyltransferase, a SAM-dependent methyltransferase, and the previously discussed RNA cap guanine-N2 methyltransferase. The possibility of a CpV-BQ1 encoded R-M system was investigated but is unlikely as we could not identify any type II site-specific restriction enzymes in the CpV-BQ1 genome.

mA6-MTase methylates the amino group at the C6 position of adenine in specific nucleotide motifs, creating an N^6^-methyl-adenine modification (72,74). N^6^-methyl-adenine modifications are rare amongst eukaryotes but are widespread amongst *Phycodnaviridae* family members (67,72,75,76). The CpV-BQ1 genome is AT-rich, with a 67.68% AT content, which makes adenine a highly prevalent nucleotide and robust candidate for methylation. Therefore this enzyme likely specifically methylates the CpV-BQ1 genome to provide protection from degradation by nucleases.

mC5-MTase methylates the 5th carbon on the pyrimidine ring of cytosine in specific nucleotide motifs, creating a 5-methyl-cytosine modification (77,78). 5-methyl-cytosine modifications are commonly found in eukaryotic genomes and are implicated in various functions such as regulating gene expression, transposon silencing, genomic imprinting, and development (72). This modification is also found in some virus genomes including *Phycodnaviridae* family members and is thought to provide the viral genome protection from nucleases (66,72,76). The mC5-MTase encoded by CpV-BQ1 is likely required for methylation of its own genome. However, it also could be involved in manipulating its host’s cellular processes, such as downregulating the expression of host genes. To explore the CpV-BQ1 genome methylation landscape single-molecule real-time sequencing could be used to identify both N^6^-methyl-adenine and 5-methyl-cytosine modifications and gain deeper insight into the protection and gene regulation mechanisms provided by these MTases (72,79).

FkbM methyltransferase is a S-adenosyl-L-methionine (SAM) transferase dependent methyltransferase enzyme with O-methylation activity (80). FkbM was originally isolated from bacteria and has been shown to perform post-modification of the macrocyclic polyketides FK506, FK520, and the antibiotic rapamycin (81). FkbM has also been found in *Phycodnaviridae* genomes, however, its substrate interactions and function is unknown (66,67,82). The final methyltransferase encoded by CpV-BQ1 could not be assigned a specific function and is designated a general SAM-dependent methyltransferase (83). SAM-MTases are a broad group of methyltransferases found in all domains of life which serve many different biological functions, thus, the functional role of this CpV-BQ1 encoded enzyme is unknown (84).

### Sugar Manipulation

Three glycosyltransferase proteins were identified in the CpV-BQ1 genome. Glycosyltransferases attach sugar moieties to proteins, which can constitute important post-translational modifications. Glycosidic bonds are catalyzed using sugar donors which contain either a nucleoside phosphate or a lipid phosphate leaving group (85,86). Glycosyltransferases are classified into hierarchical groups based on families, clans, and fold structures (87). A total of 137 glycosyltransferase (GT) families have been classified to date, listed on the carbohydrate-active enzymes (CAZy) database (http://www.cazy.org/) (88). Of the three identified GTs in the CpV-BQ1 genome, two were classified as members of specific families, while the third could not be classified as a specific GT family member.

The GT encoded at locus BQ1_30 belongs to the family GT2. The GT2 family is one of the largest GT groups and contains members with a wide variety of catalytic activities. These enzymatic functions include, but are not limited to, cellulose synthase, chitin synthase, hyaluronan synthase, and β-glucosyltransferase (87). GT2 family members contain a GT-A fold defined by two closely positioned β/α/β Rossmann domains, and use an inverting mechanism during catalysis of the donor substrate (86).

The GT encoded at locus BQ1_106 belongs to the family GT17, a small GT group in which all family members exhibit β-1,4-N-Acetylglucosaminyltransferase (GnTIII) activity and use an inverting mechanism during catalysis (87). In vertebrates, GnTIII catalyzes the formation of bisecting *N-acetylglucosamine* (GlcNAc) residues on *N*-glycans within the Golgi apparatus. This modification inhibits the action of branching enzymes, preventing the formation of highly branched *N*-glycan structures (89,90). In viruses, glycans are commonly *N*-linked to a glycoprotein asparagine residue via GlcNAc (91). These modifications are prevalent on glycoproteins in virion envelopes and play important roles in viral infection stages, including progeny formation and cellular infection (91).

Moreover, some NCLDV virions have glycans attached to the surface of major capsid proteins via the activity of GTs (92). Thus, possible functions of the three GTs encoded by CpV-BQ1 may be the synthesis of *N*-glycans on major capsid proteins or glycoproteins in the virion envelope, however, these possibilities have yet to be explored experimentally.

### Protein and lipid binding, synthesis, and modifications

The CpV-BQ1 genome encodes two proteins that likely participate in protein synthesis, the translation initiation factor 4E (eTIF4E) and DNAJ. The eTIF4E facilitates binding of the host’s ribosome to the 5’ prime cap of mRNA which initiates protein translation (93). DNAJ, originally identified in prokaryotes, functions as a cochaperone protein to Hsp70 and aids in protein folding (94). However, studies of DNAJ homologs in viruses show this protein has diverse roles and can also be involved in genome replication, transcriptional activation, virion assembly, and cellular transformation (94). Thus, further study is required to determine the exact functional nature of the DNAJ protein encoded by CpV-BQ1. The CpV-BQ1 genome also encodes three tRNAs, including tRNA-Leu, tRNA-Arg, and tRNA-Ile, which are required for incorporation of these three amino acids into nascent peptide chains (44).

Protein modification enzymes encoded by CpV-BQ1 indicate the host ubiquitination pathway is utilized by this virus to modulate proteins and perhaps subvert host defense mechanisms. Ubiquitination of proteins can either modulate their activity or signal their degradation (51). Four E3 ubiquitin ligase proteins are encoded by the CpV-BQ1 genome. Interestingly, one of the E3-ligases encoded by the genome was denoted as a N1r/p28-like protein. The N1r/p28 protein was the first E3-ligase discovered in *Poxviridae*, it is recruited to virus factories within the cell and is an important virulence factor (51,95). Thus, the CpV-BQ1 encoded N1R/p28 protein may be involved in the ubiquitination of important proteins within virus factories. Other proteins which may be involved in protein modification include six F-box domain containing proteins, which are thought to be involved in the ubiquitin-ligase complex (96), as well as, two proteins containing a metallopeptidase (WLM) domain, which are also thought to be associated with ubiquitin-signaling pathways (97).

Lastly, the patatin-like phospholipase protein encoded by CpV-BQ1 has also been found in many other NCLDVs however its function is not well understood (98). In plants, patatin phospholipase catalyzes the cleavage of fatty lipids from membranes, while in bacteria, this protein has been implicated in the pathogen-host interaction (99,100).

### DNA Packaging and Genome Completeness

The CpV-BQ1 genome encodes the viral A32 protein found in all NCLDVs which is essential for the packaging of viral DNA into virions (98,101). Silencing of the viral A32 protein results in virion structures devoid of viral DNA (102). A32 is thought to form a hexameric ring on the membrane surface of immature virions and pumps complete viral DNA into the virion (50).

In addition to the A32 protein, packaging of many NCLDV genomes, including Poxviridae, ASFV, Phycodnaviridae, and Mimiviruses, requires the presence of inverted terminal repeats on the distal ends of their genomes (59,103,104). These inverted repeats contain, in order, genes, tandem DNA repeats, and mismatched hairpin ends which interact with packaging enzymes ensuring complete genomes are incorporated into virions (50). The CpV-BQ1 genome has inverted repeats 3928 bp long at its distal ends which contain identifiable genes and tandem repeats. The possibility of hairpin sequences in the first 30, 40, 50, and 100 bp of the genome was investigated using the DNA secondary structure prediction tool by vector builder (105) (Figure 7). Of the four analyzed sequences, the 50 bp sequence provides the most promising mismatched hairpin structure, as it is similar to the stem-loop organization of the vaccinia virus hairpin motifs (106).

**Figure 7.**
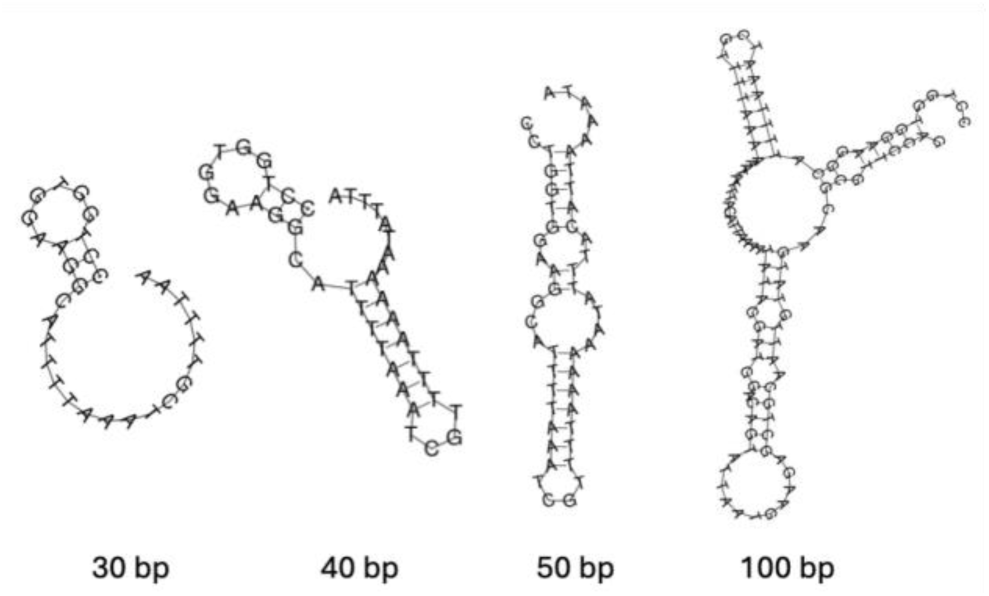
Potential Hairpin Structures of CpV-BQ1 Genome Ends. Possible secondary DNA hairpin structures formed by the 30, 40, 50, and 100 bp ends of the CpV-BQ1 genome modeled using the DNA secondary structure tool by VectorBuilder (105).

The presence and structure of mismatched hairpin ends can not be verified by genome assembly alone. However, based on the presence of terminal distal repeats and their well-defined role in other NCLDVs as well as preliminary secondary structural analysis, it is likely that the CpV-BQ1 genome forms hairpin structures. Most importantly, since distal inverted repeats are a signature of many NCLDV genome ends, the presence of these inverted repeats on the distal ends of the CpV-BQ1 genome provide strong evidence that the CpV-BQ1 genome has been sequenced completely.

### Virion capsid and associated structural proteins

Four putative viral capsid proteins were identified in the CpV-BQ1 genome. InterPro analysis of putative capsid proteins at locus BQ1_139, BQ1_140, and BQ1_141 identified an adenovirus hexon domain in these three proteins. The adenovirus hexon domain contains a jelly-roll fold which is a signature feature of NCLDV capsid proteins (107,108). Furthermore, these three proteins and BQ1_142 had a high degree of structurally similarity to other NCLDV capsid structures when analyzed with the homology-based protein fold recognition tool Phyre2 (109). Lastly, these proteins are encoded sequentially on the same strand. This proximity would permit the synthesis of all four genes in tandem, making their synthesis highly efficient. NCLDV capsid proteins have a high degree of sequence variation; a protein Blast analysis of these four proteins did not detect any sequence homology with other proteins (59). Thus, if these putative capsid proteins are indeed structural components of the virion capsid, this would expand our knowledge of NCLDV capsid sequences and aid in the identification of other capsids. Additionally, two minor capsid proteins were identified in the CpV-BQ1 genome. The minor capsid P9 transmembrane helices containing protein has domains similar to the inner capsid P9 structure of the NCLDV PBCV-1 (108). The second minor protein is a tape measure protein, which likely spans from one virion isohedral vertex to another and is thought to play a critical role in mediating the capsid size and maintaining the orientation of capsomers to one another during assembly (110).

The CpV-BQ1 genome also encodes two tail fiber proteins, one of which contains a fibronectin III binding domain. Tail fiber proteins are found in bacteriophages and facilitate the injection of viral DNA into cells. However, they have also been found in some NCLDVs including in *chloroviruses* and other *Phycodnaviruses* (67,111). In PBCV-1, tail fiber proteins form a spike on a vertice of the icosahedral capsid structure which interacts with the host cell to facilitate virion entry (111,112). Similarly, the CpV-BQ1 genome tail fiber proteins may form a spike on the capsid to facilitate cellular entry.

An important post-translational NCLDV capsid protein modification for assembly and stabilization is the formation of disulfide bonds between conserved cysteine residues (51,113). These bonds are catalyzed by a redox reaction requiring two enzymes, a thioredoxin domain containing protein and an ERV sulfhydryl oxidase (69,113). The CpV-BQ1 genome encodes an ERV/ALR sulfhydryl oxidase and two thioredoxin domain containing proteins, a thioredoxin and a glutaredoxin-like protein. DeepLoc subcellular predictions identified ER signaling domains on the ERV/ALR sulfhydryl oxidase and glutaredoxin-like protein. Together, these two proteins likely complete the redox reaction necessary for assembly and stabilization of capsid structures. DeepLoc predicted the thioredoxin protein is within the cytoplasmic compartment; however, this does not definitively exclude it from involvement in the capsid redox cascade reaction. Lastly, the CpV-BQ1 genome encodes an Ac78 gene which is similar to the baculovirus protein Ac78 (114). This protein is important for budded virion production, embedding of virions into occlusion bodies, and primary cellular infection. It has been shown to play an important role in virion localization however is not essential for virion assembly or structure (114).

### Comparison of CpV-BQ1 and CpV-BQ2 Genomes

With the publication of this work we now have the full genome sequence of two viruses, CpV-BQ1 (this study) and CpV-BQ2 (13), which can infect *C. parva*. CpV-BQ1 and CpV-BQ2 are taxonomically assigned to the NCLDV families *Phycodnaviridae* and *Mesomimiviridae* respectively. One striking difference between CpV-BQ1 and CpV-BQ2 is their genome size. CpV-BQ1 has a 165,454 bp genome with 32.32% GC content that encodes 193 ORFs, whereas CpV-BQ2 has a 437,255 bp genome with 25% GC content that encodes for 503 ORFs (13). This difference in both size and coding potential indicates that although they share the same host, they likely use remarkably different infection and replication strategies. For example, CpV-BQ2 encodes at least eight restriction-modification (R-M) systems and 13 methyltransferases (13), whereas CpV-BQ1 does not encode any R-M systems and only encodes 4 methyltransferases. This indicates these viruses use very different methods to protect their own DNA and/or degrade host DNA which can alter cellular processes such as metabolism, transcription, and translation (72,73). Furthermore, the presence of an RNA polymerase (RNA pol) II gene in CpV-BQ2 and its absence in CpV-BQ1 indicate these viruses use very different infection and replication strategies. RNA pol encoding viruses can transcribe their own DNA, NCLDVs that encode their own RNA pol typically carry out transcription, genome replication, and virion packaging all within the host’s cytoplasm (55). On the other hand, NCLDVs that do not encode an RNA pol require a nuclear infection step to hijack their host’s transcriptional machinery which is followed by a cytoplasmic infection wherein virion packaging occurs (56). Thus, CpV-BQ1 likely utilizes an two-step infection/replication strategy that includes both cytoplasmic and nuclear infection which requires coordination across subcellular compartments, whereas CpV-BQ2 likely infects *C. parva* through a one-step cytoplasmic approach.

## Conclusions

In this study we have sequenced and annotated the complete linear 165,454 bp genome of Chrysochromulina parva virus BQ1 (CpV-BQ1). CpV-BQ1 was originally isolated from a lake in Ontario and is a lytic agent of the haptophyte alga *Chrysochromulina parva.* Taxonomic analysis of polB, A32 ATPase, and VLTF-3 protein sequences were used to assign the *Phycodnaviridae* family classification to CpV-BQ1. The genome contains 193 genes, of which 92 could be assigned a known function. CpV-BQ1 has hallmark genes found in many NCLDVs necessary for genome replication, virion production, and transcription (3). Like other phycodnaviruses, CpV-BQ1 most likely has a two-step cellular infection life-cycle, first entering the nucleus for transcription as evidenced by the lack of an RNA polymerase gene, then virion assembly ensues within the cytoplasm (56,64). Previously, another *C. parva* lytic agent Chrysochromulina parva virus BQ2 (CpV-BQ2) was isolated from the same water sample as CpV-BQ1 (11), its genome was sequenced and was taxonomically assigned to the *Mesomimiviridae* family (13). Thus far, co-infection of a eukaryotic algae with viruses from both *Phycodnaviridae* and *Mesomimivirdae* families has only been observed in the species *Phaeocystis globosa* (115,116). However, only the complete genome sequence of one *P. globosa* infecting virus, the group II PgV-16T mimivirus, has been sequenced and made available (117). It is postulated that co-infections of eukaryotic algae with viruses belonging to different NCLDV families is common, however, due to a lack of data this hypothesis cannot be currently supported (115). Indeed, only ∼60 eukaryotic algal viruses have been isolated in culture (115), while thousands of eukaryotic algal species have been isolated and are available in culture collections around the world. This disparity emphasizes the lack of research and knowledge surrounding eukaryotic algal virus diversity, taxonomy, life cycle, and environmental impact. With the work reported here, we have for the first time established an algal-virus system with complete genome sequences for both *Phycodnaviridae* and *Mesomimiviridae* viruses. This system can be used to study the biological, ecological, and environmental consequences of the coding potential of these viruses which replicate in the same host. For example, the presence of RM systems encoded by CpV-BQ2 and their absence in CpV-BQ1 may be relevant to inter-viral competition and suggest that BQ2 may restrict the replication of BQ1, or the PLVs which presumably parasitize one of these viruses. With the genomic information in hand, detailed transcriptional studies can be conducted to further understand the complicated dynamics and relationships between *Chrysochromulina parva* and its viral parasites. In turn, this knowledge will illuminate the potential complexities of algal virus-host interactions and is especially important considering the critical importance of algae in the biosphere and human affairs.

## Author Contributions

D.N. conducted the molecular work, Nanopore sequencing, genome assembly and annotation, plotted the data and interpreted initial results, and wrote the original draft of the manuscript; C.P. and I.I. conducted virus culturing and purification, extraction of virus nucleic acids, and Illumina sequencing; T.C.C., J.I.N. and SMS acquired funding, supervised and administered the project; J.I.N. and S.M.S. conceived the work, interpreted results, and co-wrote the manuscript. All authors revised, edited, and approved the final manuscript.

## Acknowledgments

This study was funded by a Natural Sciences and Engineering Research Council of Canada (NSERC) Discovery Grants (2022-03066) awarded to S.M.S; NSERC Discovery Grants (2022-03350 and 2022-00329) awarded to J.I.N.; and Mitacs Accelerate funding (IT18981) awarded to T.C.C and D.N.

## Data Availability

The CpV-BQ1 genome sequence is available on GenBank under accession PQ783904.

Raw reads are available on the SRA database BioProject accession number PRJNA1199504. BioSample accessions for raw reads are as follows: Illumina whole genome sequences, SAMN45676274; Nanopore whole genome sequences, SAMN45876275; CpV-BQ1 PCR amplified region 1, SAMN45876276; CpV-BQ1 PCR amplified region 2, SAMN45876277; CpV-BQ1 PCR amplified region 3, SAMN45876278; CpV-BQ1 PCR amplified region 4, SAMN45876279; CpV-BQ1 PCR amplified region 5, SAMN45876280.

