## Supplementary figures and images for "Hybrid sequencing reveals the genome of a *Chrysochromulina parva* virus and highlight its distinct replication strategy"

### Supplementary Table 1

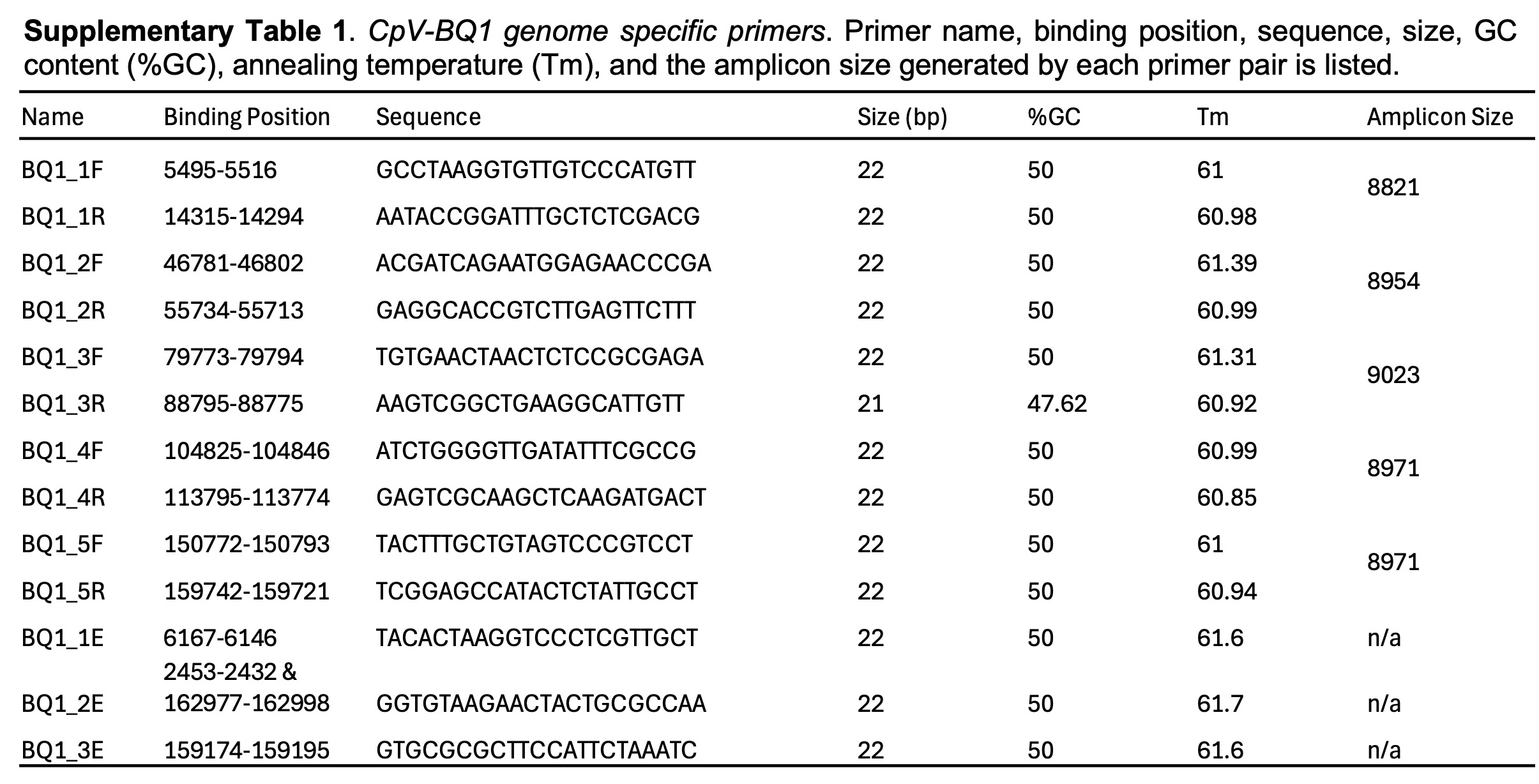

### Supplementary Table 2

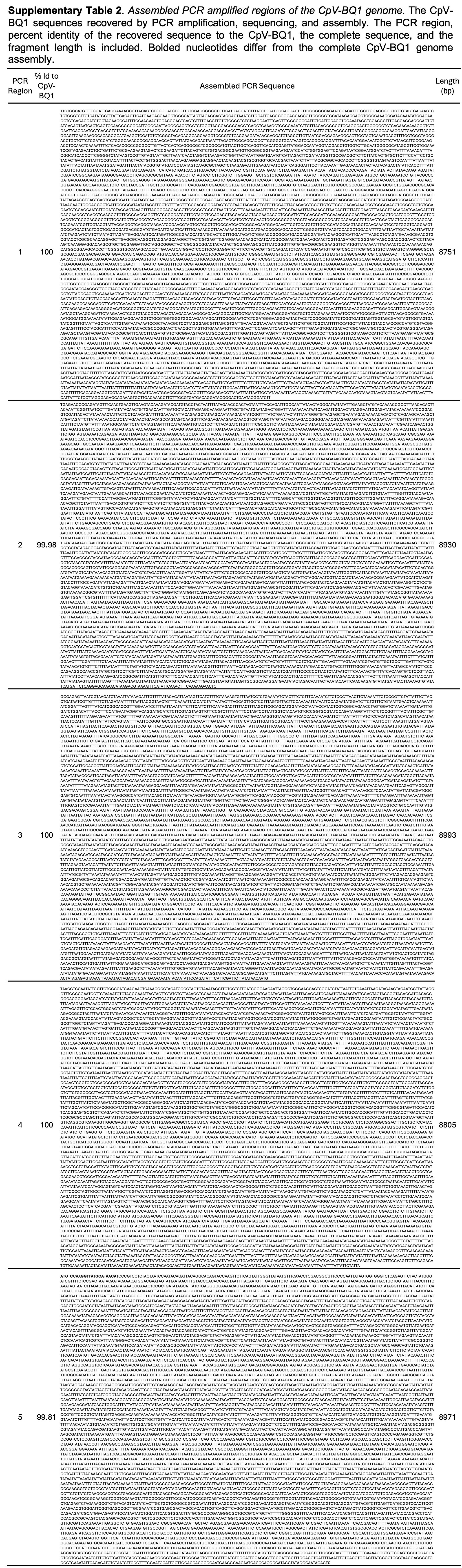

### Supplementary Table 3

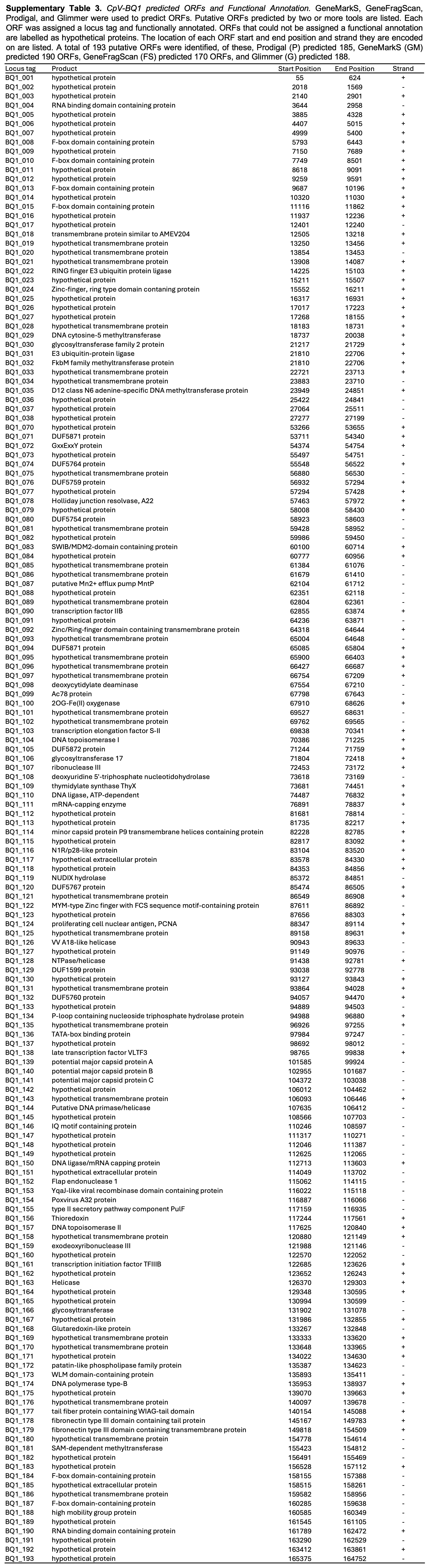

### Supplementary Table 4

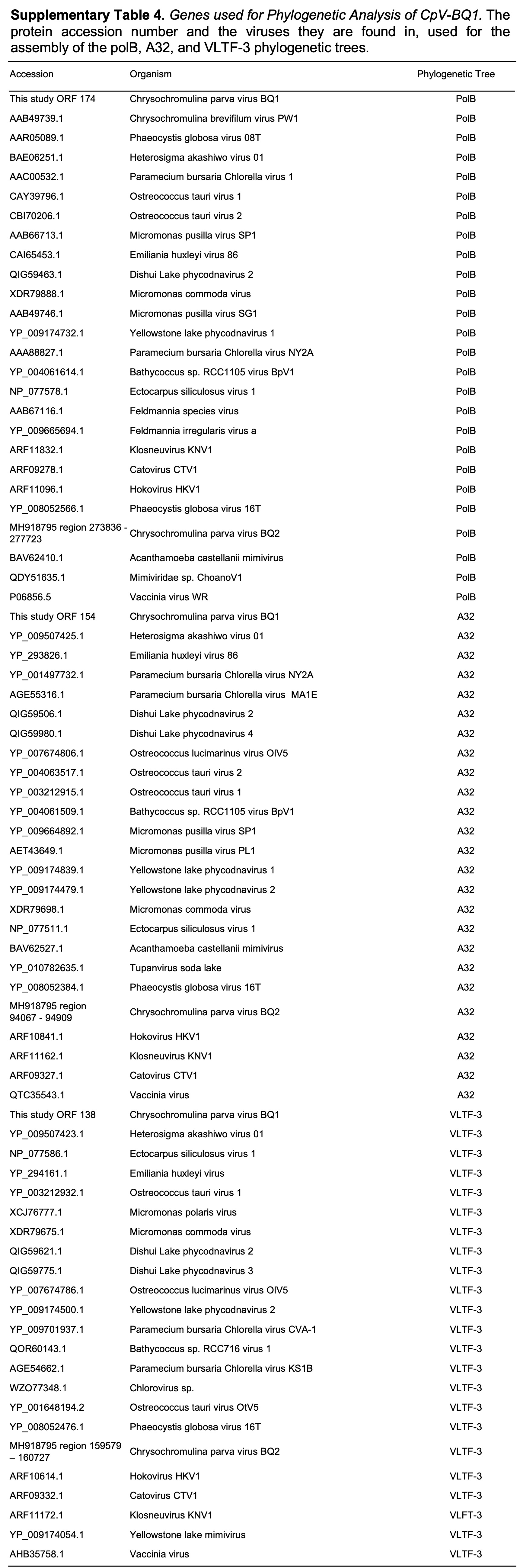
